# Transcriptomic analysis of melanin production in *Exophiala dermatitidis* conditional albino mutants on two different carbon sources

**DOI:** 10.1101/2025.09.23.676125

**Authors:** Kamaldeep Chhoker, Georg Hausner, Steven D. Harris

**Affiliations:** Department of Biological Sciences, University of Manitoba, Winnipeg, MB, Canada; Department of Microbiology, University of Manitoba, Winnipeg, MB, Canada; Department of Plant Pathology, Entomology and Microbiology, Iowa State University, Ames, IA, USA

**Author notes:** Corresponding author Department of Plant Pathology, Entomology and Microbiology, Iowa State University, 1344 ATRB, 2213 Pammel Dr., Ames, IA 50014 USA.

**Keywords:** *Exophiala dermatitidis*, 1, 8-DHN melanin, DOPA melanin, Pyomelanin, PKS1, Conditional albino, Obligate albino, RNA-transcriptomics

## Abstract

*Exophiala dermatitidis* is a polymorphic black yeast found in various habitats and man-made environments such as soil, sinks and saunas. Melanin plays a key role in *E. dermatitidis* virulence and environmental adaptation. The *E. dermatitidis* genome sequence has revealed the presence of genes responsible for production of melanin via three different pathways (1,8-DHN melanin, DOPA melanin, and pyomelanin) but besides DHN melanin not much is known about the activation of the other pathways. Our previous work identified three conditional albino mutants (*alb1*, *alb2* and *alb3*) that can recover melanin production despite mutation in *PKS1*. In this study RNA Transcriptomics was used as a tool to investigate gene expression differences between the three conditional albinos, two obligate albinos (*alb10* and *alb12*) and wildtype to account for variation in melanization using two treatments (dextrose and galactose) as carbon sources. Differential gene expression analysis revealed a higher number of significantly upregulated genes in the conditional albinos on galactose (YPG) compared to the obligate albinos. Overrepresentation analysis revealed a higher number of significantly enriched GO terms in the conditional albinos compared to the wildtype and the obligate albinos. Multiple genes involved in 1,8-DHN melanin and pyomelanin were also found to be upregulated on YPG in the conditional albinos. Besides genes involved in melanization, genes involved in various aspects of cell wall regulation were also upregulated on YPG. To this date this is the first study that has demonstrated the activation of genes involved in multiple different melanin pathways in *E. dermatitidis* grown on different carbon sources.

## Introduction

Fungal melanin plays a wide variety of roles that include protection from harmful environmental conditions (Salgado-Castillo et al. 2023) as well as the promotion of virulence in fungal pathogens of plants, animals and humans (Okuno *et al*. 1983; Howard and Valent 1996; Dean 1997; Adachi *et al*. 1998; Schnitzler et al. 1999;) . The main function attributed to fungal melanin is protection from UV irradiation (Patel et al. 1996; Khajo et al. 2011), which in some cases is associated with the quantity of melanin produced by cells (Dighton et al. 2008; Cordero and Casadevall 2017). Other studies have also demonstrated the ability of fungal melanin to confer protections by binding to toxic metals such as copper (Gadd and de Rome 1988), iron (Saiz-Jiminez and Shafizadeh 1984), zinc and tin (Rizzo et al. 1992).

*Exophiala dermatitidis* also known as *Wangiella dermatitidis* is an ascomycete black yeast with a world-wide distribution that can be found in natural habitats such as tropical rain forests, where it is most likely associated with fruits and wild fruit-eating animals (Sudhadham et al. 2008). There has also been resurgence of *E. dermatitidis* recovered from man-made environments such as bathrooms, kitchen sink drains, saunas, steam baths and railways ties (Matos et al. 2002; Hamada and Abe 2010). The emergence of *E. dermatitidis* as an opportunistic pathogen has been noted in immunocompromised patients where it is able to cause phaeohyphomycosis, chromoblastomycosis and fatal infections of central nervous system (de Hoog 2014; Olsowski et al. 2018; Vasquez et al. 2018; Hagiya et al. 2019; Beniwal et al.2023). Several studies have identified DHN-melanin synthesized via the polyketide synthase Pks1 as a key attribute related to virulence in *E. dermatitidis*. Notably, mutant strains lacking Pks1 and therefore unable to produce melanin show reduced virulence on skin and mouse model infections (Chen et al. 2014; Poynter et al. 2016; 2018). Genome annotation of *E. dermatitidis* has revealed that Besides 1,8-DHN melanin, genome annotation has revealed that *E. dermatitidis* also possesses the capacity to produce L-dihydroxyphenylalanine (L-DOPA) melanin and pyomelanin via the L-tyrosine degradation pathway (Chen *et al*. 2014; Poyntner *et al*. 2018; Carr *et al*. 2023).

As part of the ongoing study on the genetics of melanin production in *E. dermatitidis*, our previous work employed classical and unbiased random mutagenesis approach to generate albino mutant strains (Chhoker et al. 2025b). Unexpectedly, all recovered albino mutants possess mutations in the *PKS1* gene. The *alb* mutant strains could further be subdivided into conditional and obligate depending on the ability of different carbon sources to restore melanin production (Chhoker et al. 2025b). Specifically, the study identified 17 albino mutants all possessing a mutation in *PKS1*, but a subset of these, termed conditional albino (*alb1*, *alb2*, and *alb3*), were able to restore pigmentation on YPG. The ability of conditional albinos to restore pigmentation highlights the fact that fungal species that have genes involved in multiple different melanin biosynthetic pathways might be able to restore pigmentation if certain conditions are met.

Previous studies on albino mutants of *E. dermatitidis* obtained via targeted mutations of *PKS1* have never observed pigment restoration, making this study the first of its kind where albino mutants of *E. dermatitidis* could restore pigmentation presumably via utilizing alternative melanin biosynthetic pathways. We propose that comparison of gene expression profiles in the conditional and obligate *alb* mutants presents an opportunity to identify genes and pathways involved in the activation of these alternative pathways, thereby providing novel insight into the regulation of melanin production in the polyextremotolerant fungus *E. dermatitidis*.

## Materials and Methods

### Media and strains

Two different media were selected for this study: Yeast extract peptone dextrose (YPD) and Yeast extract peptone galactose (YPG) (Table 1)

**Table 1:**
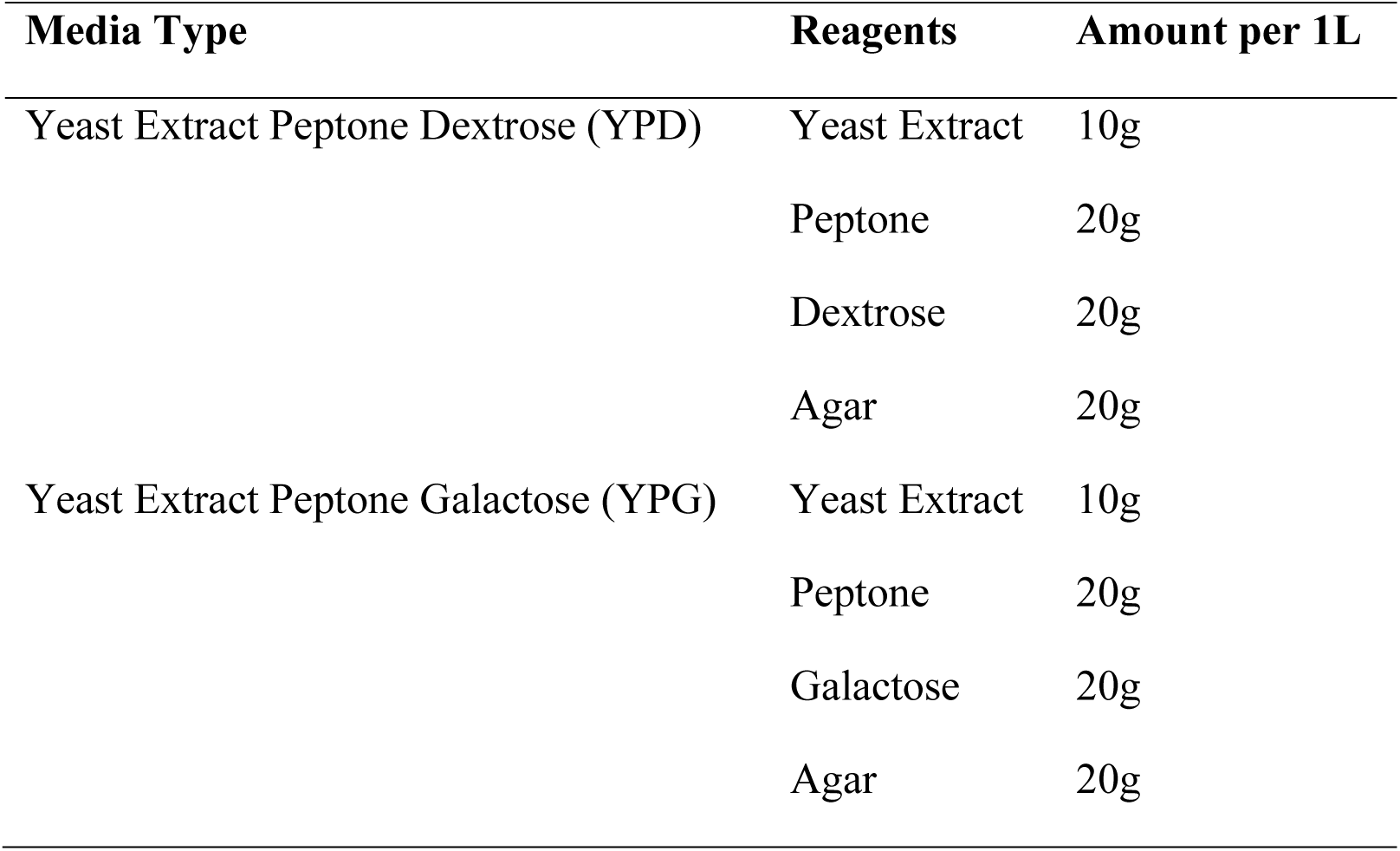
Different media types and recipes used in this study.

| Media Type | Reagents | Amount per 1L |
| --- | --- | --- |
| Yeast Extract Peptone Dextrose (YPD) | Yeast Extract | 10g |
|  | Peptone | 20g |
|  | Dextrose | 20g |
|  | Agar | 20g |
| Yeast Extract Peptone Galactose (YPG) | Yeast Extract | 10g |
|  | Peptone | 20g |
|  | Galactose | 20g |
|  | Agar | 20g |

### Fungal strains

The *Exophiala dermatitidis* reference strain UT8656 (=ATCC 34100, = CBS 525.76) was treated as the wildtype strain as described in Chen et al. (2014). Five *alb* mutants including three conditional albinos (*alb1*, *alb2*, and *alb3*) which produce melanin on YPG but not on YPD and two obligate albinos (*alb10* and *alb12*) which have completely lost their ability to produce melanin on both YPD and YPG, were selected for this study (Chhoker et al. 2025b) (Figure 1).

**Figure 1:**
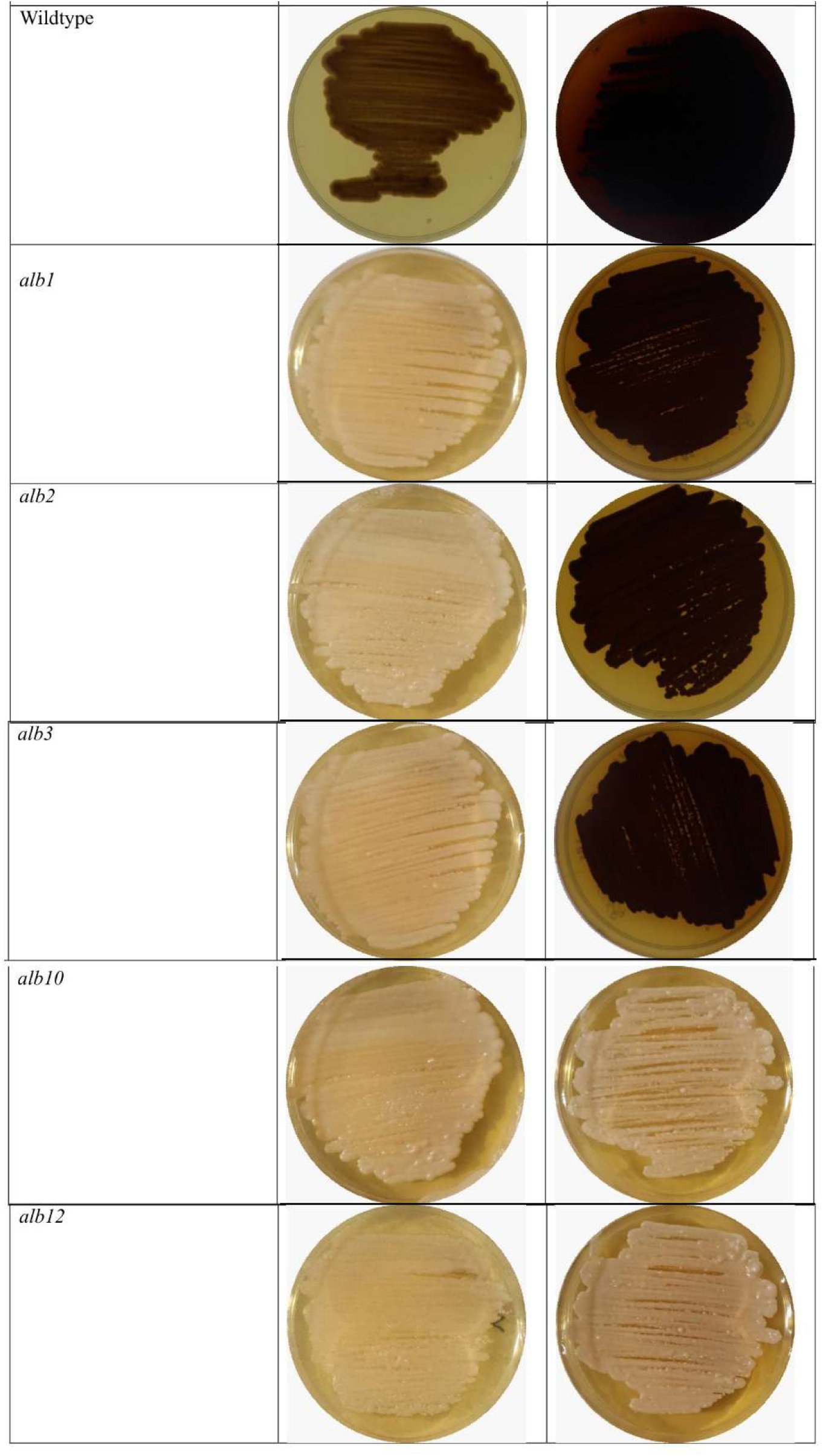
Images representing various strains of *E. dermatitidis* growth after 7 days at 28°C. Growth observed on YPD on the left and YPG on the right.

Strains were obtained from −80°C stocks stored in 30% glycerol and grown at 28°C for a week until colonies started to appear on YPD plates. Single colonies were obtained, plated on YPD or YPG plates, and grown at 28°C for 7 days. Three replicates for each strain were used to obtain a total of 18 YPD plates and 18 YPG plates giving us 36 samples in total. After 7 days of growth, the fungal strains were subjected to RNA extraction.

### RNA extraction and Sequencing

Total RNA from all 36 samples was extracted using the RNeasy Mini Kit (Qiagen, Germantown, MD, United States) following manufacturers guidelines with slight modifications. Briefly, material from seven-day old YPD and YPG plates was scraped off using sterile flat toothpicks and added to 2ml screw cap collection tubes containing 500µl RLT buffer and 5µl β-ME (β-mercaptoethanol) on ice. Silica beads were added, and the tubes were run on a Bead Beater (Sigma-Aldrich, Missouri, United States) for 15 seconds to homogenize the samples.

Samples were maintained on ice to minimize degradation, and all centrifugation was performed for 20 seconds at 8000G. DNA contamination was removed using the RNase-Free DNase Kit (Qiagen) following manufacturers’ guidelines.

Initial readings to check the quality and quantity of extracted RNA were obtained using a NanoDrop 2000/2000c Spectrophotometer (Fisher Scientific). The Agilent TapeStation 4150 (AgilentTechnologies, Santa Clara, CA, United States) was used to obtain RIN values. Any sample that yielded RNA with a RIN value below 7 was re-extracted to obtain higher molecular weight RNA. The RNA (Minimal concentration >30 (ng/μL) was shipped to the GenomeQuebec Innovation Centre (McGill University, Quebec, Canada). The RNA samples used for library preparations were enriched for PolyA mRNA, and sequencing of the RNA libraries (cDNA) was performed in both directions (paired-ends reads of 100bp) on an Illumina NovaSeq 6000 platform (PE100 4 X 25 million reads/lane) aiming for 100million reads per sample.

### Transcriptomic analysis

RNA-seq analysis was performed on data obtained from all 36 samples utilizing tools available on the Galaxy platform (https://galaxyproject.org/usegalaxy/) (The Galaxy Community 2024). First, Fastq files containing the raw reads (as data obtained from GenomeQuebec) were subjected to quality checks using FastQC version 0.12.1 (Andrews 2010). Low Quality bases with a phred score below 20 and adaptor sequences were trimmed using cutadapt version 5.9 (Martin 2011). The cleaned and trimmed sequences were mapped against the *E. dermatitidis* reference genome sequence (NCBI Refseq: GCF_000230625.1) using HISAT2 version 2.2.1 with paired end reads (Kim et al. 2015). The BAM files were visualized using the IGB browser (Freese *et* al. 2016). MultiQC version 1.27 (Ewels et al. 2016) was used after each step to summarize the analysis results from various tools to visualize all samples at once. FeatureCounts version 2.0.6 (Liao et al. 2013) was used to count the number of mapped reads to annotation elements present in the reference genome. The count matrix obtained using FeatureCounts for three replicates per treatment (YPD vs YPG) was used to identify differentially expressed genes (DEGs) using DESEQ2 version 2.11.40.8 (Love et al. 2014). Genes were considered differentially expressed between the two conditions (YPD vs YPG) if their adjusted p-value (referred to as FDR) was <0.05. Volcano plots were constructed to observe the number of significant up and down regulated genes based on FDR <0.05 and Log2(FC) > 2 or < −2. The obtained lists of differentially expressed genes for each of the *alb* mutants was compared with the lists of differentially expressed genes of the wildtype using BioVenn (https://www.biovenn.nl/index.php) (Hulsen et al. 2008) and were analysed for enriched Gene Ontology (GO terms) using FungiFun3 (https://fungifun3.hki-jena.de/) (Priebe et al. 2014; Lopez et al. 2024). The Fisher exact test was used to calculate the significance of the enriched GO terms and Benjamini-Hochberg Correction was used to obtain adjusted p-values.

## Results

### Overview of differentially expressed genes (DEGs)

The number of DEGs varied between the wildtype and the five mutant *alb* strains when observed using a significance cutoff of *p* < 0.05 and Log2(FC) > 2 or <-2 (Supplemental Figure S1). Several trends were observed. First, the numbers of upregulated and downregulated genes were greater in the albino mutants compared to wildtype, with the exception of the alb12 mutant that had fewer upregulated genes than wildtype (Figure 1). Second, all the conditional albinos (*alb1*, *alb2* and *alb3*) had a higher number of differentially upregulated genes (>400) and a higher number of downregulated genes (>1000) compared to the obligate albinos (*alb10*, and *alb12*) that had <400 upregulated genes and <1000 downregulated genes (Figure 2). Magnitude of down-regulation was greater for all albino mutants and the wildtype. And, lastly, the number of shared upregulated and downregulated genes between the mutants and the wildtype were quite low (Figure 2).

**Figure 2:**
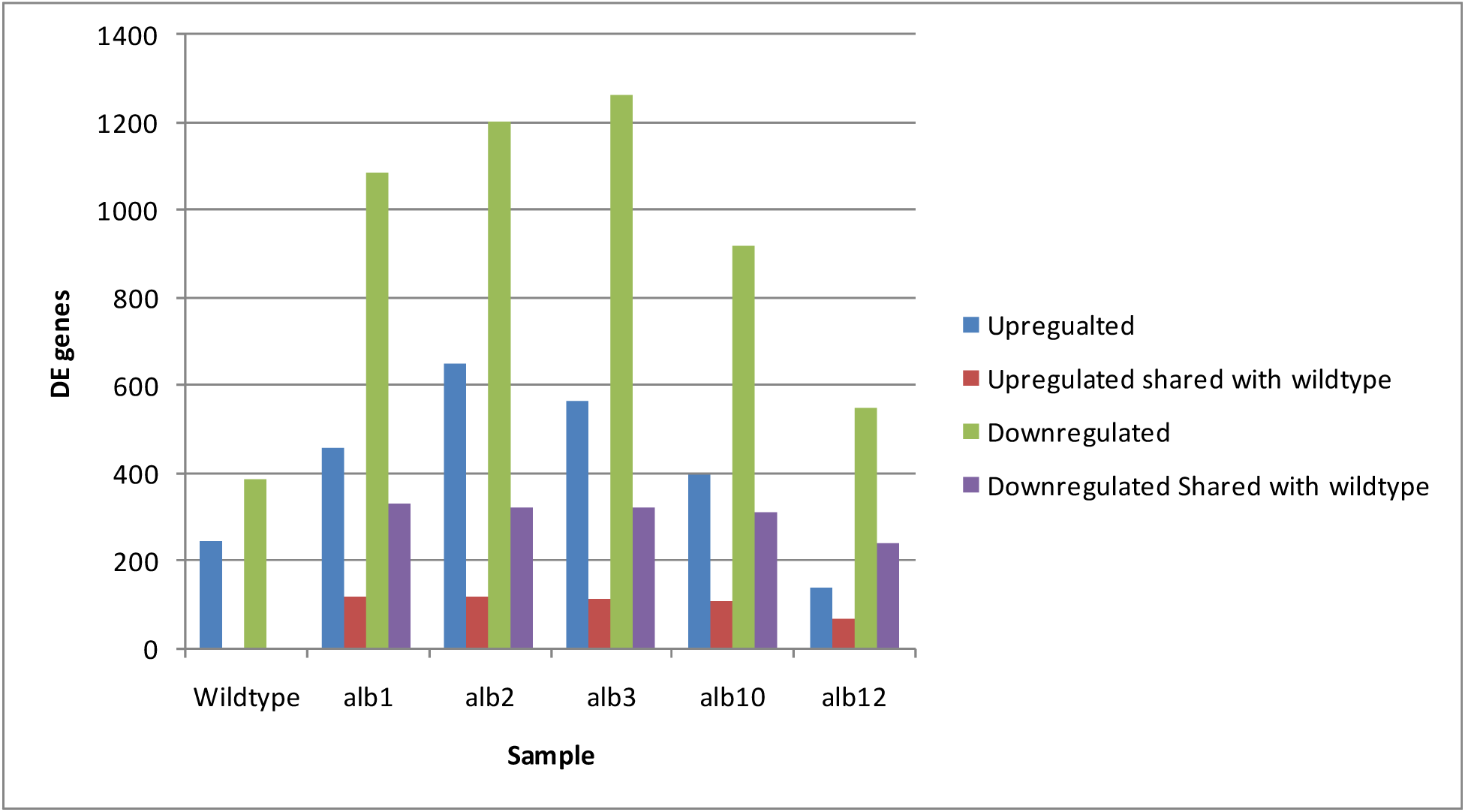
The number of genes differentially expressed, with a significance cutoff of *p* <0.05 and Log2(FC) > 2 or < −2 for each mutant strain, as well as shared with the wildtype. Graph represents the number of DEGs using the count matrix data obtained using FeatureCount with three replicates for each per treatment (YPD vs YPG).

### Overrepresentation Analysis

To determine which GO terms were enriched among the various samples, Overrepresentation Analysis (ORA) (Chicco and Agapito 2022) was performed using the FungiFun3 platform. For each mutant strain, Venn diagrams represent gene enrichment analyses based on unique and shared differentially expressed upregulated and downregulated genes between the wildtype and mutant *alb* strains (Supplemental Figure S2). Based on the ORA analysis performed on differentially expressed genes (DEGs) with *p <* 0.05 and Log2(FC) >2 or <-2, some trends appeared. With regards to shared upregulated genes between the wildtype and the *alb* mutants, none of the mutants showed any significantly enriched GO terms at FDR cutoff of 0.05 (Supplemental Figure S2 a-e). However, the wildtype strain showed significant enrichment in microtubule binding, microtubule motor activity and microtubule-based movement when considering upregulated genes not shared with the *alb* mutants (Supplemental Figure S2 a-e). The conditional *alb* mutants displayed a variable number (ranging from nine to twelve) of significantly enriched GO terms when subjected to one-by-one comparison with wildtype (Supplemental Figures S2 a-c). Both obligate albino mutants *alb10* and *alb12* had the lowest number with just one enriched GO terms (Supplemental Figure S2 d, e). Strikingly, there were no shared upregulated GO terms amongst all the *alb* mutants.

Overrepresentation analysis on significantly expressed unique downregulated genes revealed that, the wildtype strain did not show any enriched GO terms for unique genes when compared with *alb1*, *alb2*, *alb3* and *alb10* mutants but had three enriched GO terms (obsolete oxidation-reduction process, transaminase activity and oxidoreductase activity while comparing with *alb12* (Supplemental Figure S2 f -j). With regards to shared downregulated genes between the wildtype and the *alb* mutants, conditional albino mutants *alb1*, *alb2* and *alb3* had five, four and three enriched GO terms respectively, whereas obligate albino *alb10* had six enriched GO terms and *alb12* had the highest number with 14 enriched GO terms (Supplemental Figure S2 f-j). The conditional *alb* mutants displayed a variable number (ranging from one to three) of significantly enriched unique GO terms when subjected to one-by-one comparison with wildtype (Supplemental Figures S2 f-j). Obligate albino mutant *alb10* had no unique enriched GO terms and *alb12* had the highest number with 13 unique enriched GO terms when subjected to one-by-one comparison with the wildtype (Supplemental Figure S2 f-j). There were no shared downregulated GO terms amongst all the *alb* mutants.

### Shared GO terms between albino mutants

Amongst up-regulated genes, there were four significantly enriched GO terms shared between all conditional *alb* mutants, transmembrane transport, transmembrane transporter activity, plasma membrane and zinc ion binding. Notably, none of these terms were found in the obligate albino mutants. One GO term (DNA-binding transcription factor activity, RNA polymerase II-specific) was only shared between the *alb1* and *alb2* conditional albinos. Two GO terms (acyl-CoA dehydrogenase activity and acylformamidase activity) were shared between the *alb2* and *alb3* conditional albinos (Figure 3). The only significantly enriched GO term that was found in both conditional and obligate albino mutants was DNA-templated transcription which was enriched in all three conditional albinos (*alb1*, *alb2*, and *alb3*) and in one obligate albino (*alb10*) (Figure 3).

**Figure 3:**
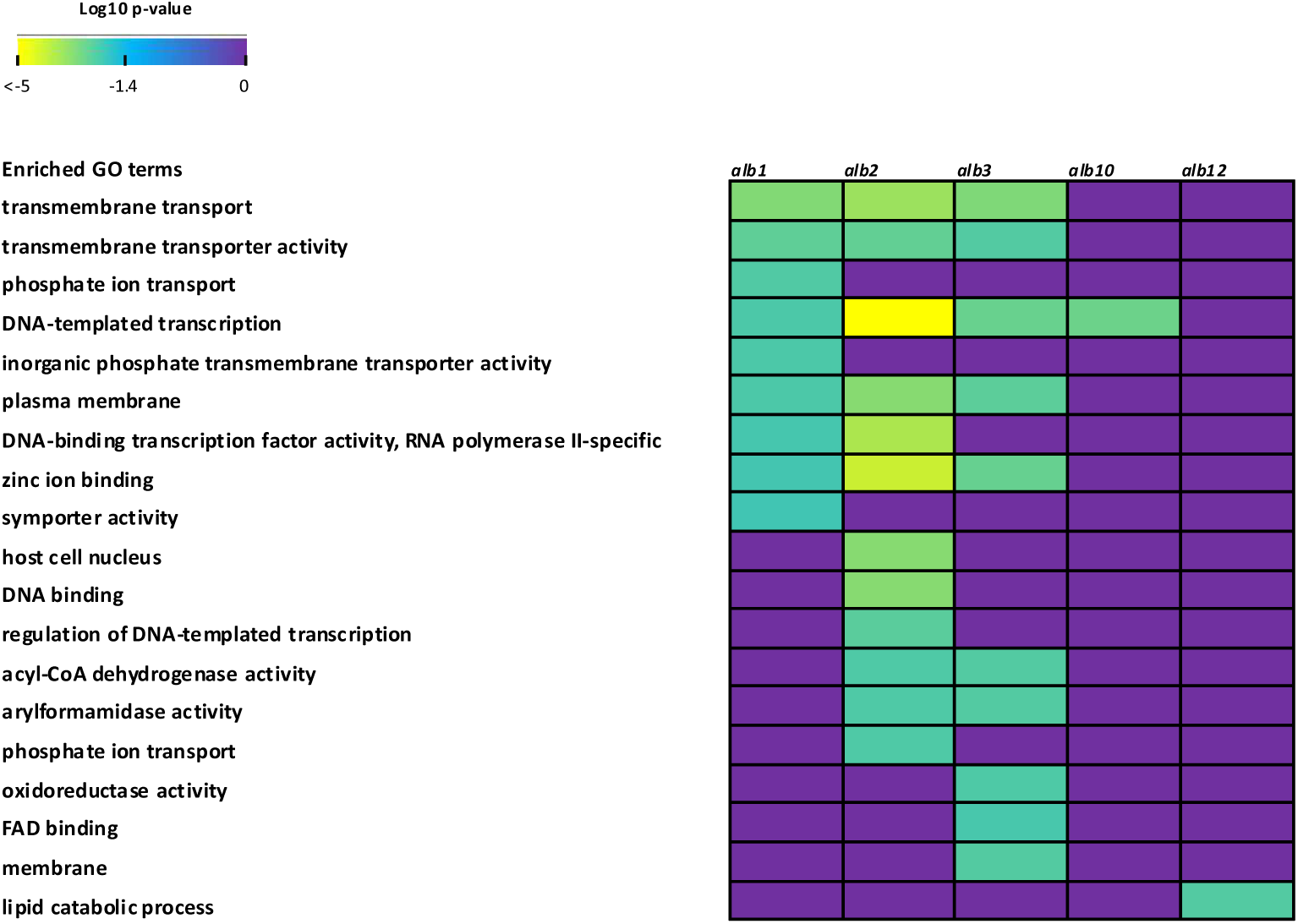
Heatmap representing the significantly enriched GO terms (FDR < 0.05) obtained from the FungiFun 3.0 platform that are shared between the five albino mutants based on differentially expressed upregulated genes with *p* < 0.05 and Log2FC >2. Yellow colour = more statistically enriched, purple colour= less statistically enriched.

Most of the genes involved in the GO term transmembrane transport and transmembrane transporter activity were related to major facilitator superfamily transporters (MFS transporter). Other genes in these two GO terms corresponded with retrograde regulation protein and amino acid protein transporters. The GO term plasma membrane consisted of genes involved in MFS transporters, multiple genes involved in chitin synthesis and a Ras2 homologue. The GO term zinc-ion binding encompassed genes encoding various transcription factors (TFs) and transcriptional regulators such as GAL4 involved in galactose metabolism and multiple protein kinases such as stk16 protein kinase homologue (Supplemental File 1).

Based on downregulated DEGs with *p* < 0.05 and Log2(FC) <-2, there were no GO terms that were significantly enriched amongst all the albino mutants (Figure 4). However, the GO term carbon-sulfur lyase activity was shared between all conditional albino mutants (*alb1*, *alb2*, and *alb3*). This term encompasses genes encoding glutathione-dependent formaldehyde (GFA) activating enzymes (Supplemental File 1). An additional GO term transmembrane transport was significantly enriched among the conditional albino mutants *alb2* and *alb3* (Figure 4).

**Figure 4:**
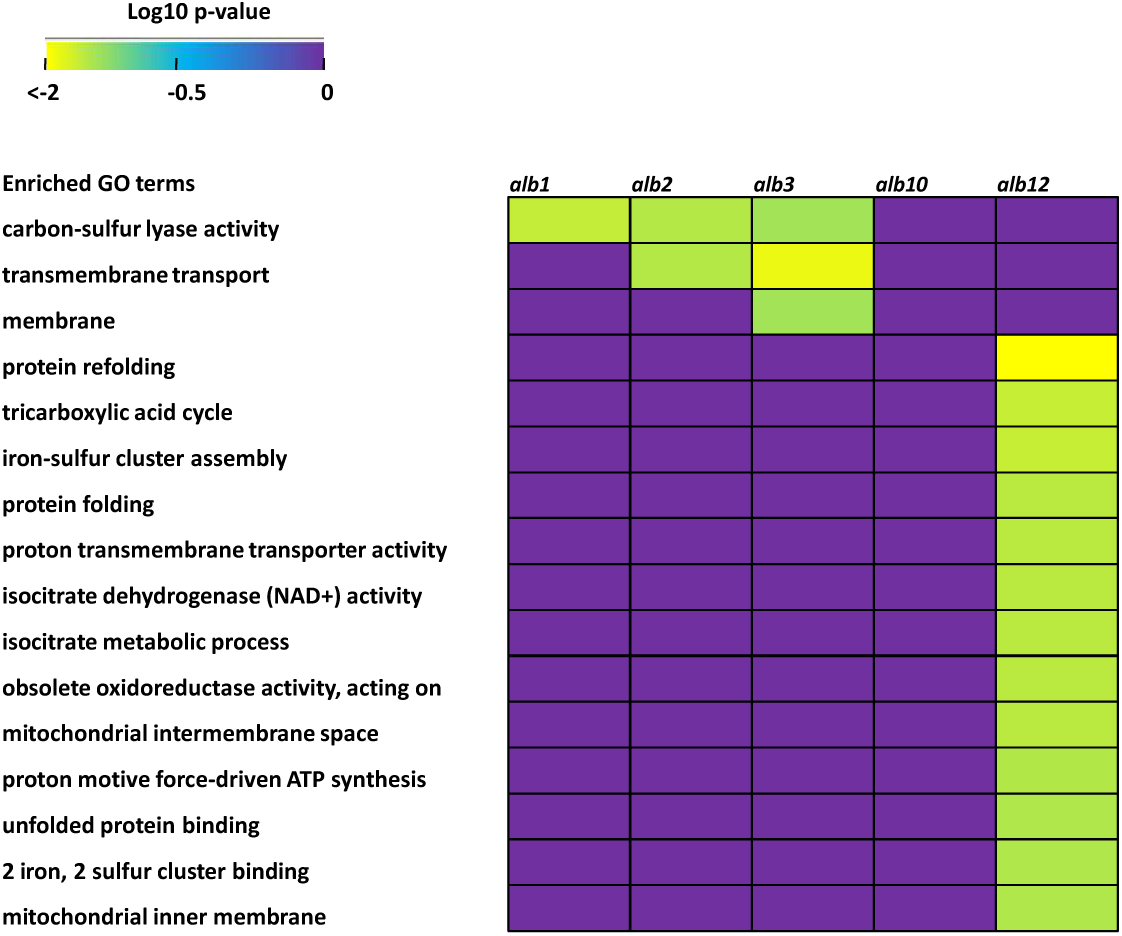
Heatmap representing the significantly enriched GO terms (FDR < 0.05) obtained from the FungiFun 3.0 platform that are shared between the five albino mutants based on differentially expressed downregulated genes with *p* > 0.05 and Log2FC < −2. Yellow colour = more statistically enriched, purple colour= less statistically enriched.

### Melanin biosynthetic pathways

Based on DEGs obtained via DESEQ2 between the two treatments (YPD vs YPG), there were several genes involved in the synthesis of 1,8-DHN melanin that were found to be upregulated (Figure 5). The HMPREF1120_03173 (Polyketide synthase), HMPREF1120_05939 (1,3,6,8-Tetrahydroxynaphthalene reductase), HMPREF1120_07724 (Scytalone dehydratase) and HMPREF1120_02828 (Fungal pigment MCO) genes were each significantly upregulated in all three conditional albinos (*alb1*, *alb2*, and *alb3*) and the obligate albino *alb10*.

**Figure 5:**
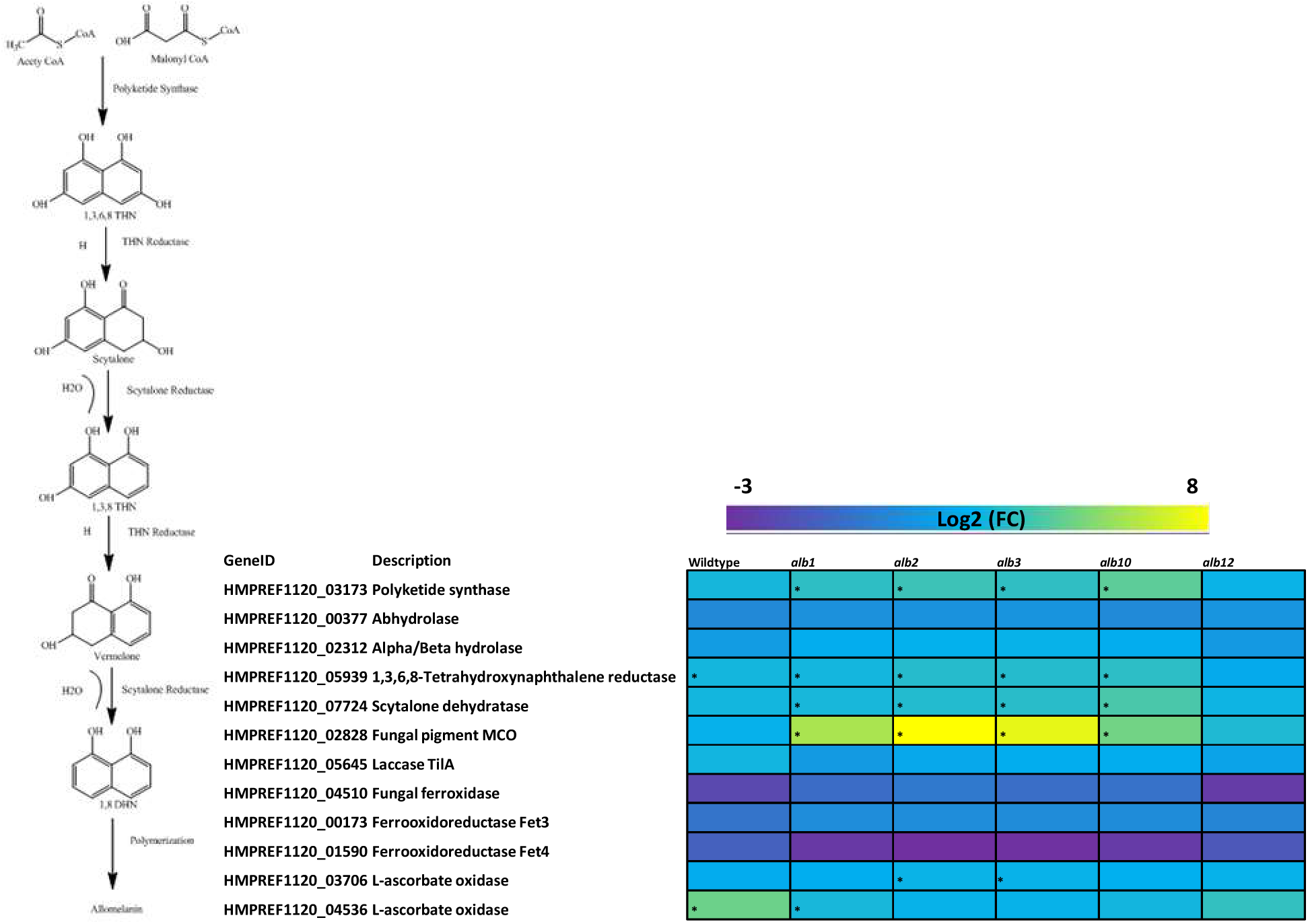
Heatmap representing the DEGs involved in the 1,8-DHN melanin biosynthetic pathway found to be upregulated or downregulated based on two treatments (YPD vs YPG) and three replicates each for the wildtype and three conditional albinos (*alb1*, *alb2* and *alb3*) and two obligate albinos (*alb10* and *alb12*). Yellow colour = upregulated DEGs, purple colour= downregulated DEGs. * Indicate significantly upregulated DEGs (Log2FC >1 and *p <0.05*).

HMPREF1120_03706 (L-ascorbate oxidase) was only upregulated in two conditional albinos (*alb2* and *alb3*) and HMPREF1120_04536 (L-ascorbate oxidase) was upregulated in the wildtype and conditional albino *alb1* (Figure 5).

For the tyrosine degradation pathway, several genes were found to be upregulated on YPG. None of the genes involved in the tyrosine degradation pathway were upregulated in the wildtype and the obligate albino *alb12* (Figure 6). HMPREF1120_02164 (Tyrosine aminotransferase) was significantly upregulated in all three conditional albino *alb1*, *alb2* and *alb3* and in the obligate albino mutant *alb10*. HMPREF1120_05584 (4-Hydroxyphenylpyruvate dioxygenase) was significantly upregulated in conditional albinos *alb2* and *alb3*, and obligate albino *alb10*. HMPREF1120_03825 (Fumarylacetoacetate hydrolase) was significantly upregulated in only one conditional albino *alb2*. HMPREF1120_03438 (Maleylacetoacetate isomerase) was significantly upregulated in all three conditional albinos (*alb1*, *alb2* and *alb3*), and obligate albino *alb10*. One gene, HMPREF1120_03827 (Homogentisate dioxygenase) was significantly downregulated in the wildtype, conditional albinos *alb3* and the obligate albino alb10 (Figure 6).

**Figure 6:**
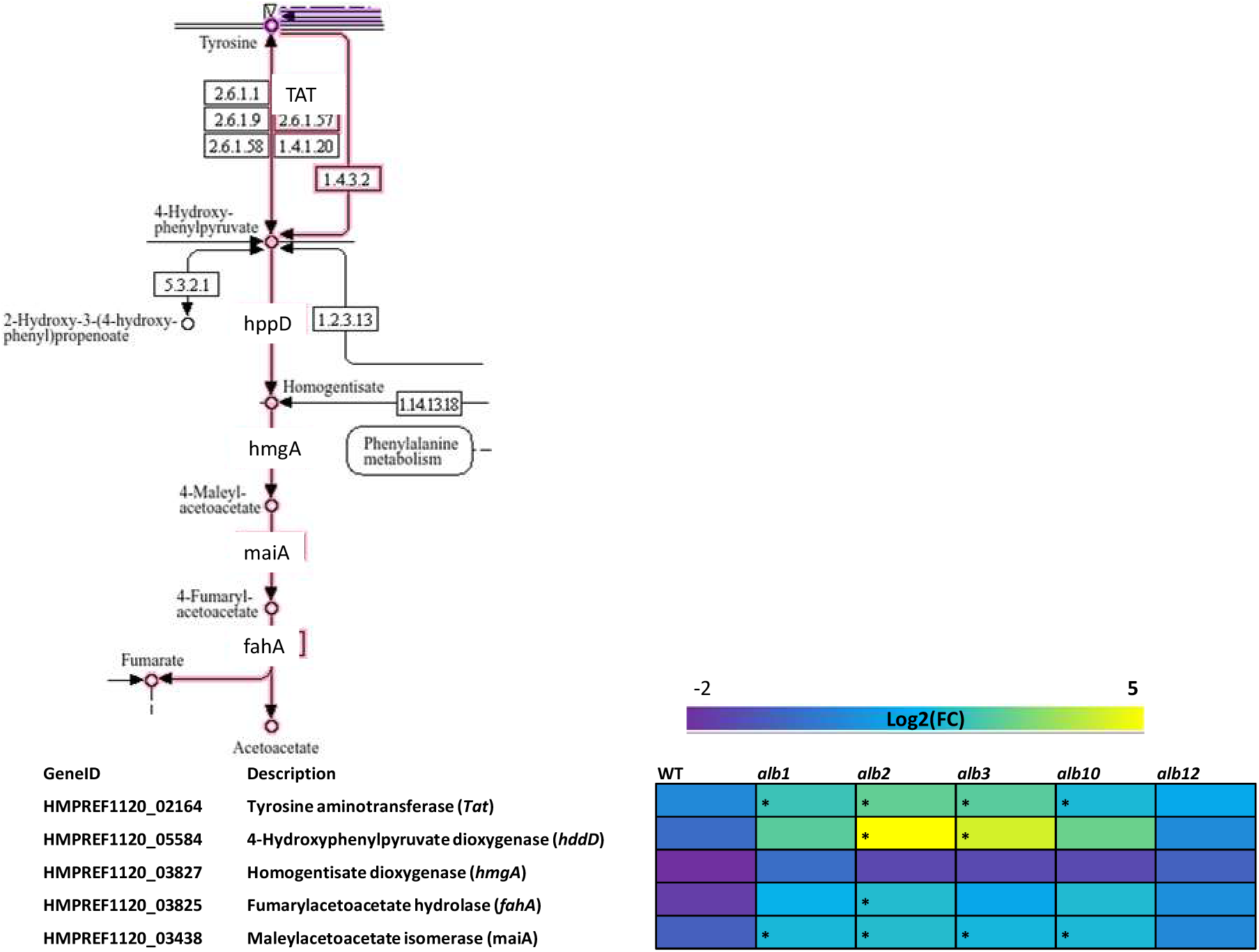
Heatmap representing the DEGs involved in the tyrosine degradation pathway found to be upregulated or downregulated based on two treatments (YPD vs YPG) and three replicates each for the wildtype and three conditional albinos (*alb1*, *alb2* and *alb3*) and two obligate albinos (*alb10* and *alb12*). Yellow colour = upregulated DEGs, purple colour= downregulated DEGs. * Indicate significantly upregulated DEGs (Log2FC >1 and *p <0.05*).

Most of the genes involved in the DOPA melanin pathway did not show any upregulation in response to different treatments (YPD vs YPG) (Figure 7). HMPREF1120_07692 (Tyrosinase) showed significant upregulation in the wildtype and obligate albino *alb12*. Another gene, HMPREF1120_00199 (Multicopper oxidase) was found to be upregulated in the wildtype and in all albino mutants (*alb1*, *alb2*, *alb3*, *alb10* and *alb12*) (Figure 7).

**Figure 7:**
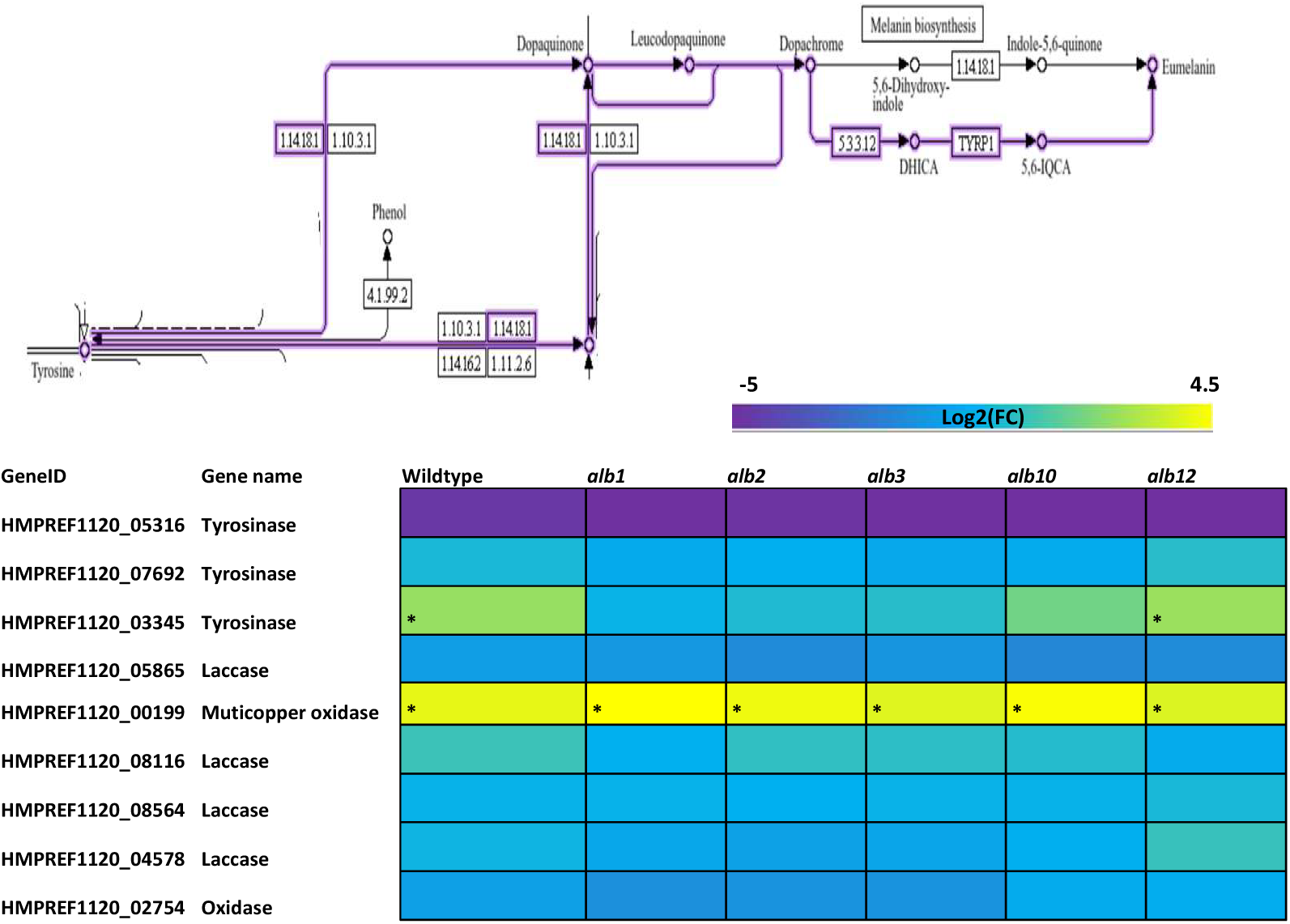
Heatmap representing the DEGs involved in the DOPA melanin pathway found to be upregulated or downregulated based on two treatments (YPD vs YPG) and three replicates each for the wildtype and three conditional albinos (*alb1*, *alb2* and *alb3*) and two obligate albinos (*alb10* and *alb12*). Yellow colour = upregulated DEGs, purple colour= downregulated DEGs. * Indicate significantly enriched upregulated DEGs (Log2FC >1 and *p <0.05*).

### Secondary metabolism and light sensing

Based on the DESEQ2 analysis between the two treatments (YPD vs YPG), a small number of genes involved in the production of secondary metabolites appear to be significantly upregulated on YPG (Figure 8a). HMPREF1120_06543 (Superkiller protein 3) and HMPREF1120_08091 (3-oxoacyl-[acyl-carrier-protein] synthase II) were significantly upregulated in all three conditional albinos (*alb1*, *alb3* and *alb3*) but not in the obligate *alb* mutants or the wildtype (Figure 8a). HMPREF1120_02993 (Acetylaranotin,toxin) was upregulated in all three conditional albinos (*alb1*, *alb2* and *alb3*) and the wildtype.

**Figure 8:**
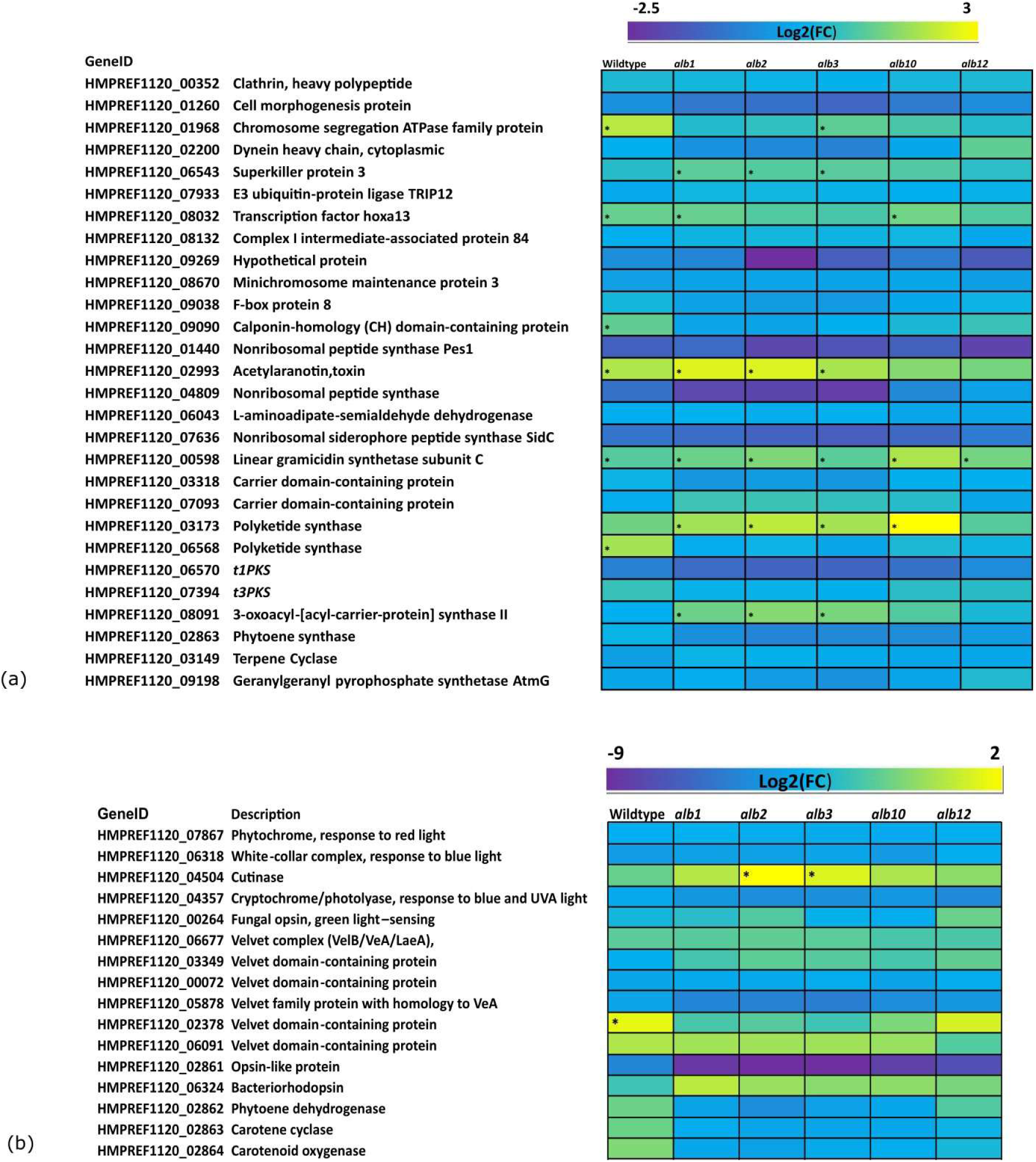
Heatmap representing the DEGs involved in (a) secondary metabolism, and (b) light sensing in *E. dermatitidis* found to be upregulated or downregulated based on two treatments (YPD vs YPG) (analysis based on three replicates for all strains examined). Significance based on DESeq2 analysis (*p* < 0.05 and Log2(FC) <1 or >-1). Yellow colour = upregulated DEGs, purple colour= downregulated DEGs. * Indicate significantly upregulated DEGs (Log2FC >1 and *p <0.05*).

HMPREF1120_01968 (Chromosome segregation ATPase family protein) was upregulated in the wildtype and the conditional albino *alb3*. HMPREF1120_08032 (Transcription factor hoxa13) was significantly upregulated in the wildtype, conditional albino *alb1* and obligate albino *alb10*. HMPREF1120_00598 (Linear gramicidin synthetase subunit C) was upregulated in the wildtype, and all five albino mutants. HMPREF1120_09090 (Calponin-homology (CH) domain-containing protein) and HMPREF1120_06568 (Polyketide synthase) was only significantly upregulated in the wildtype.

Two light sensing genes appear to be upregulated on YPG. HMPREF1120_04504 (White collar 2) was upregulated in the conditional albinos alb2 and alb3 and HMPREF1120_02378 (Velvet domain-containing protein) was only upregulated in the wildtype (Figure 8b). Most other light sensing genes did not show any upregulation but there were a few genes that were highly downregulated on YPG (Figure 8b)

### Cell wall regulation

Genes involved in the cell wall biosynthesis, were found to be upregulated on YPG (Figure 9). HMPREF1120_06816 (CHS2 Class I chitin synthase), HMPREF1120_06669 (Class V chitinase) and HMPREF1120_04557 (Chitinase) were all found to be upregulated in the wildtype and all three conditional albinos (*alb1*, *alb2*, and *alb3*). HMPREF1120_05230 (Glucans Putative exo-1,3-b-glucanase family) was only found to be upregulated in the obligate albino *alb12*. HMPREF1120_00547 (Bgl2-family of putative 1,3-b-transglucosylases) was upregulated in the wildtype, all three conditional albinos (*al1*, *alb2*, and *alb3*) and obligate albino *alb10* (Figure 9).

**Figure 9:**
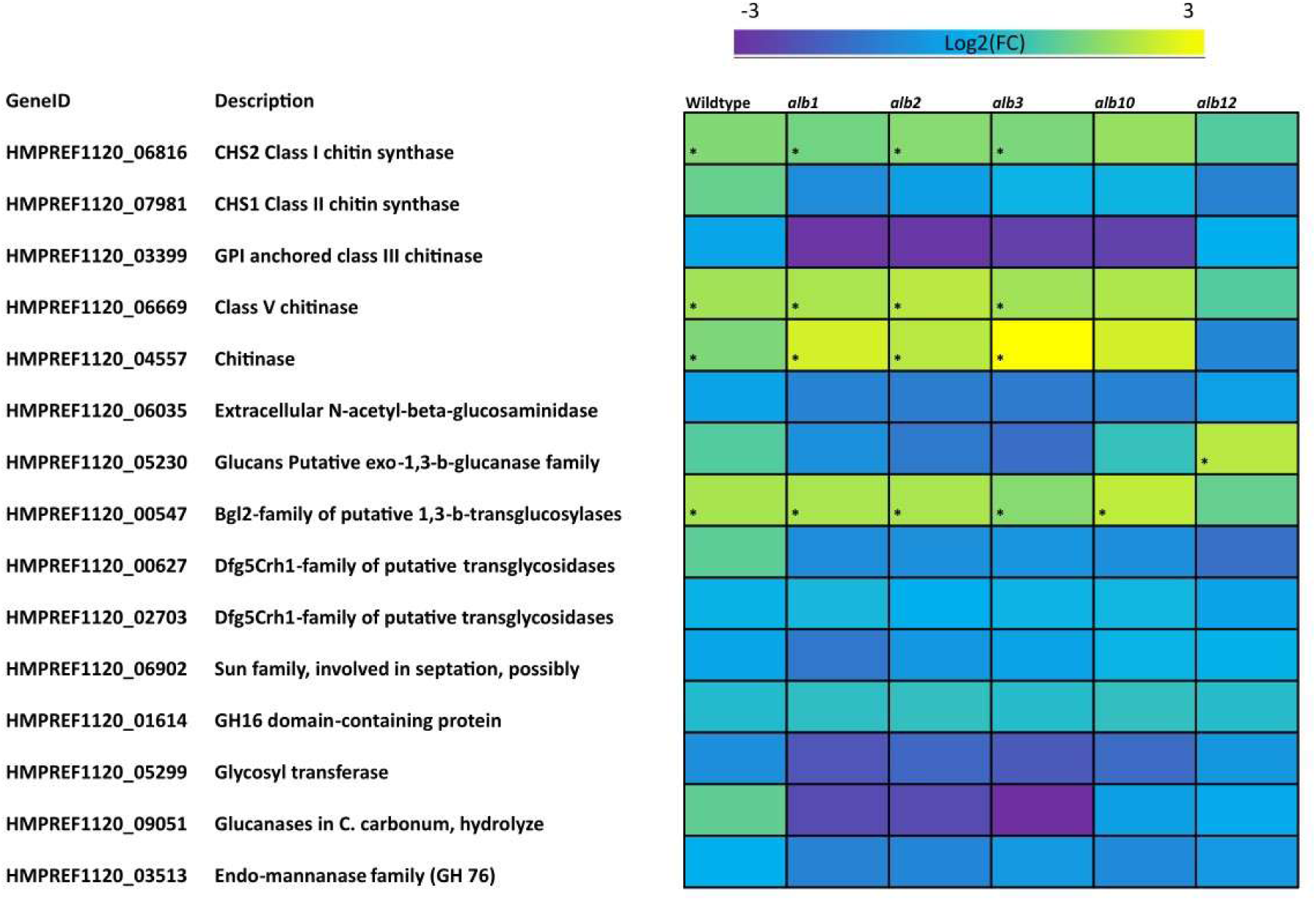
Heatmap representing the DEGs involved in cell wall regulation of *E. dermatitidis* found to be upregulated or downregulated based on two treatments (YPD vs YPG) (analysis based on three replicates for all strains examined). Significance based on DESeq2 analysis (*p* < 0.05 and Log2(FC) <1 or >-1. Yellow colour = upregulated DEGs, purple colour= downregulated DEGs. * Indicate significantly upregulated DEGs (Log2FC >1 and *p <0.05*).

### Metal acquisition

In *E. dermatitidis*, the siderophore pathway is responsible for the uptake of iron via the reduction of Fe^2+^ to Fe^3+^ (Poyntner et al. 2016). Homologues of genes encoding the enzymes involved in the siderophore pathway, namely *sidA*, *sidC*, *sidD* and *sidF* and the transporter *sit1* were all significantly downregulated in the wildtype and all albino mutants on YPG (Figure 10). *Ftr1* and *Fet3*, genes involved in the reductive uptake pathway were also significantly downregulated in the wildtype and all albino mutants on YPG. There was no significant upregulation or downregulation of genes involved in the copper transport in either the wildtype or any of the albino mutants.

**Figure 10:**
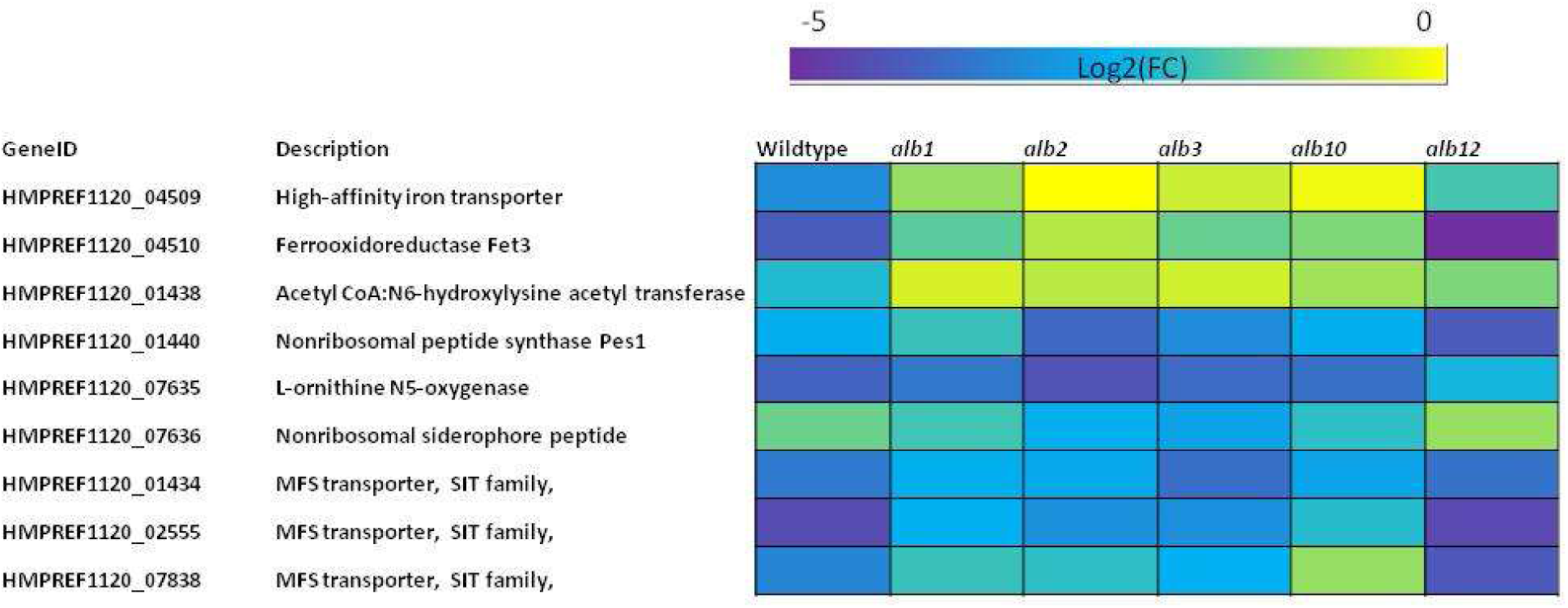
Heatmap representing the DEGs involved in metal uptake of *E. dermatitidis* found to be upregulated or downregulated based on two treatments (YPD vs YPG) (analysis based on three replicates for all strains examined). Significance based on DESeq2 analysis (*p* < 0.05 and Log2(FC) <1 or >-1. Yellow colour = slightly downregulated DEGs, purple colour= extremely downregulated DEGs.

### Carbon uptake

HMPREF1120_03751 (Aconitate hydratase), part of the glyoxylate cycle was significantly upregulated in all three conditional albinos (*alb1*, *alb2*, *alb3*) and the obligate albino mutant *alb10*. HMPREF1120_06787 (Malate dehydrogenase) was significantly downregulated in all three conditional albinos (*alb1*, *alb2* and *alb3*) and both obligate albinos *alb10* and *alb12* (Figure 11).

**Figure 11:**
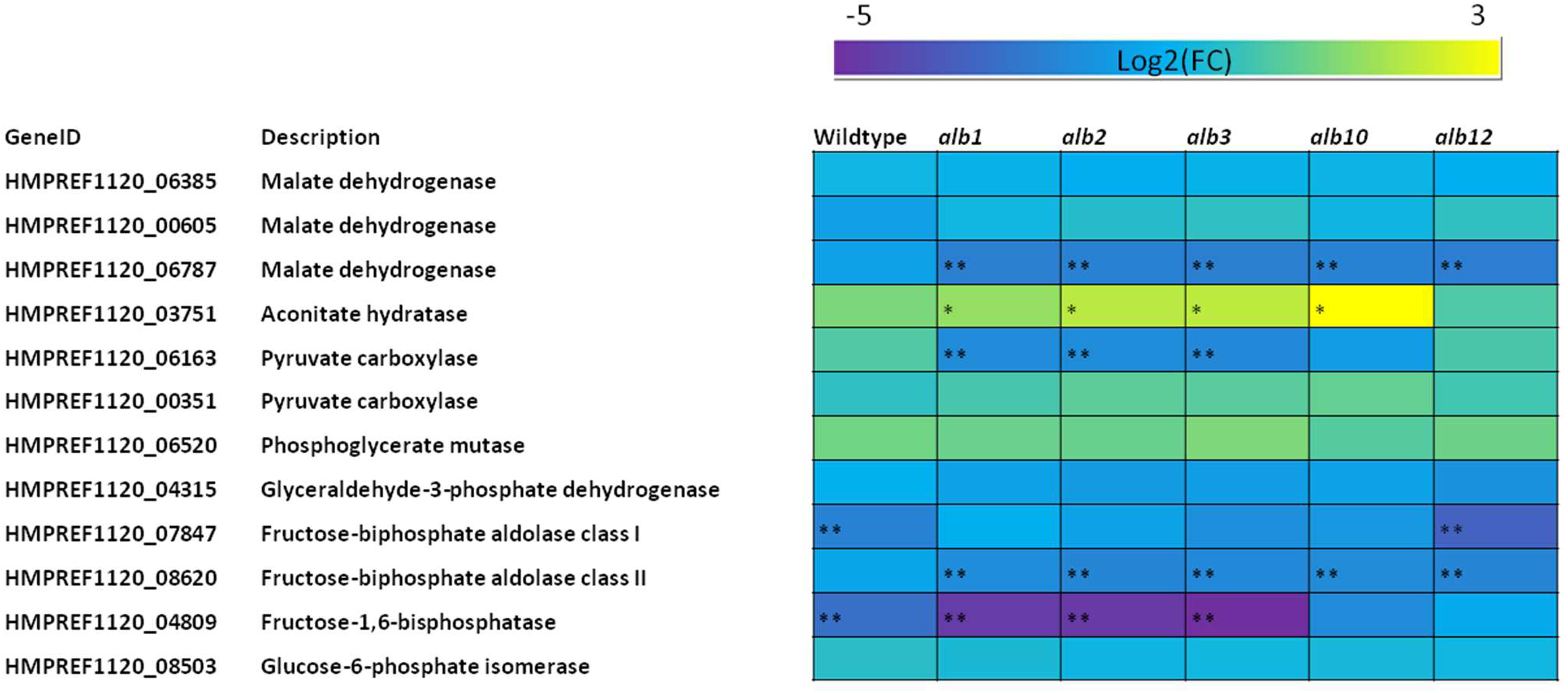
Heatmap representing the DEGs involved in the glyoxylate cycle and gluconeogenesis of *E. dermatitidis* found to be upregulated or downregulated based on two treatments (YPD vs YPG) (analysis based on three replicates for all strains examined). Significance based on DESeq2 analysis (*p* < 0.05 and Log2(FC) <1 or >-1. Yellow colour = upregulated DEGs, purple colour= downregulated DEGs. ** genes were significantly downregulated (Log2FC >1 and *p <0.05*).

In the gluconeogenesis pathway, HMPREF1120_04809 (Fructose-1,6-bisphosphatase) was significantly downregulated in the wildtype and all three conditional albinos (*alb1*, *alb2* and *alb3*). HMPREF1120_06163 (Pyruvate carboxylase) was significantly downregulated in all three conditional albinos. HMPREF1120_07847 (Fructose-bisphosphate aldolase class I) was significantly downregulated in the wildtype and obligate albino mutant *alb10*.

HMPREF1120_08620 (Fructose-biphosphate aldolase class II) was significantly downregulated in all three conditional albinos and both obligate albinos (Figure 11).

## Discussion

For this study three conditional (*alb1*, *alb2* and *alb3*) and two obligate (*alb10* and *alb12*) mutants were picked representing both categories of *alb* mutants we obtained in our previous work (Chhoker et al. 2025b). Besides mutations in *PKS1*, our previous work did not find any other mutations that were shared between all three conditional *alb* mutants, indicating that the recovery of melanization caused by the carbon source shift might be governed by multiple different pathways and mechanisms that interact with each other. This study was performed to better understand the pathways whose expression might contribute to the conditional recovery of melanin synthesis in a subset of the albino mutants. Certain trends were observed based on RNA transcriptomics. There were co-regulated genes in the wildtype and all albino mutants likely responding to the carbon shift. There were genes whose expression was shared between both the conditional and the obligate albino mutants. These are likely a response to a combination of pigment loss and the carbon shift experienced by the mutants. Lastly, there were a small number of genes that were differentially expressed in the conditional albino mutants but not in the obligate mutants, thereby potentially identifying processes that could contribute to the recovery of melanization in the conditional *alb* mutants.

### Enriched GO terms

There were no significantly enriched GO terms that were shared between all five albino mutants indicating that all five mutants behaved differently from each other. However, three GO terms (transmembrane transport, transmembrane transporter activity and plasma membrane) were only enriched amongst the conditional albino mutants. These terms each relate to membrane function and solute transport across the membrane. Most of the genes in these GO terms corresponded with MFS transporters, which play key roles in multidrug resistance and growth during stress (Chen et al. 2017). In fungi, melanin production is typically a key component of stress response, which suggests that the conditional albino mutants might have experienced some type of stress during the carbon shift that uniquely triggered melanin production. Since fungal melanization occurs in specialized vesicles that are moved and are deposited in the cell wall (Eisenman et al. 2009; Camacho et al. 2019) the shared nature of these GO terms which also included chitin synthase genes might indicate their roles in cell wall regulation which is important in melanization.

Another term that was shared between all conditional *alb* mutants was zinc ion binding. Most of the genes involved were TFs and transcriptional regulators. There is growing evidence of involvement of TFs that can act either upstream or downstream of various fungal melanization pathways to influence melanin production (Tsuji et al. 2000; Shelest 2017; Chhoker et al. 2025a). The upregulation of wide variety of TFs in the conditional albino mutants might be linked to the recovery of melanization observed in our study. Metal ions such as Zn^2+^ have been shown to result in the increase of free radical population of melanin (Buszman et al. 2006). In fungi, melanin can bind to a wide variety of metals which can also induce melanogenesis and the melanin allows for bioabsorption of essential metals from their environment (Cordero and Casadevall 2017). Since the conditional *alb* mutants were able to recover melanin production, the genes involved in bioabsorption and uptake of certain metals might be upregulated to increase the production of melanin and protect the organism from heavy metal toxicity.

### Melanization in *E. dermatitidis*

The dematiaceous nature of *E. dermatitidis* is attributed to the deposition of 1,8-dihydroxynaphthalene [1,8-DHN] melanin within the cell wall (Geis et al. 1984; Taylor *et al*. 1987; Feng et al. 2001). In *E. dermatitidis*, it has been observed that the DHN-melanin pathway genes were upregulated during pH stress, which might facilitate adaptation to its natural niches (Chen et al. 2014). In our study *PKS1* gene was upregulated in all three conditional albino mutants and one obligate albino mutant indicating that multiple genes in the PKS pathway are necessary for the production of DHN melanin. The conditional *alb* mutants in our study incorporated missense mutations (Chhoker et al. 2025b), so the *PKS1* gene might not have completely lost its function. In DHN pathway, Malonyl-CoA/acetyl-CoA acts as the precursor serving as initiator and extender units of PKS which catalyze the first step of the DHN melanin biosynthetic pathway to produce 1,3,6,8-tetrahydroxynaphthalene (T4HN) (Verde-Yáñez et al. 2023). The T4HN compound is then reduced to scytalone by THN reductase. Scytalone is dehydrated by scytalone dehydratase to form 1,3,8-trihydroxynaphthalene (T3HN). Another reduction reaction by the THN reductase forms vermelone from T3HN and the vermelone compound is then converted to DHN-melanin by scytalone dehydratase (Schumacher 2016; Verde-Yáñez et al. 2023). All three conditional albinos and one obligate albino showed an upregulation of THN reductase and Scytalone dehydrates genes, showing that parts of the DHN-melanin could be functional and possibly producing intermediates. These intermediates might then be shunted off to other processes such as fatty acid synthesis which also uses similar precursors such as acetyl-CoA and malonyl-CoA (Zhang et al. 2022). In fungi, 3-oxoacyl-[acyl-carrier-protein] synthase II enzyme is involved with the fatty acid synthesis which can be responsible for producing carotenoids in fungi (Sandmann 2022).

In fungi, pyomelanin is produced by the tyrosine degradation pathway via the oxidative polymerization of homogentisate (HGA) (Schmaler-Ripcke et al. 2008; Lorquin et al. 2022). Two enzymes: 4-hydroxyphenylpyruvic acid dioxygenase and homogentisic acid oxidase are required to degrade L-tyrosine to acetoacetate and fumarate (Ruzafa et al. 1995; Kotob et al. 1995; Schmaler-Ripcke et al. 2008). Pyomelanin is then produced from the auto-oxidation followed by self-polymerization of HGA (Fernandes et al. 2021). Upregulation of multiple genes involved in the tyrosine degradation pathway indicates that certain aspects of the pathway are being triggered in all three conditional albinos. This phenomenon has been previously observed in *E. dermatitidis* albino mutants where genes involved in the L-tyrosine pathway were upregulated during skin infection (Poynter et al. 2016). During infection, *E. dermatitidis* experiences reprogramming of its carbon metabolism, and since our treatment focused on two different carbon sources, similar phenomenon might be at play here that is inducing the L-tyrosine pathway.

Some fungi can produce pyomelanin (DOPA melanin) through a pathway that involves cysteine and DOPA quinine reactions to form various intermediates that are polymerized to generatepheomelanin. Most of the genes involved in the production of DOPA melanin did not show any upregulation in our treatments. Previous studies on *E. dermatitidis* infection did not show any upregulation of genes involved in DOPA melanin (Poynter et al. 2016) but upregulation of specific laccasses and tyronises was observed at low pH (Chen et al. 2014) indicating that specific conditions are required for the activation of the DOPA melanin pathway. Since only the carbon source was changed between the two treatments (YPD and YPG), the cells might not have experienced the required conditions to trigger the activation of the pathway.

### Secondary metabolism

In fungi, light is responsible for regulating critical aspects of life cycles including activation of biosynthetic pigments and regulation of melanin production (Tisch and Schmoll 2009; Estrada and Avalos 2008; Corrochano and Garre 2010; Dasgupta et al. 2015; Fuller et al. 2013; 2015; 2016). In *Neurospora crassa*, blue light has been shown to regulate carotenoid pigment production which is disrupted in mutants lacking the WC-1 or WC-2 photoreceptors (Froehlich et al. 2002; He et al. 2002). This has also been recorded in *Fusarium* species and *Botrytis cinerea*, where carotenoid pigment development and accumulation are regulated by light (Avalos and Estrada 2010; Canessa et al. 2013; Hevia et al. 2015). Besides melanin, *E. dermatitidis* is known to produce two carotenoids: torulene and torularhodin (Geis and Szaniszlo 1984). Since *E. dermatitidis* produces both melanin and carotenoids it can be theorized that melanin helps shield the species from light to some extent, which can then be detected by photoreceptor proteins to trigger carotenoid synthesis (Chen et al. 2014). Carotenoid production in fungi is triggered by light and most genes involved in secondary metabolism and light sensing did not show upregulation in either the conditional or obligate albinos. Since the cultures were grown in a benchtop incubator with no light:dark cycle inputs, we did not see any accumulation of carotenoids in the albino mutants, or the upregulation of genes involved in secondary metabolism. This implies that in our mutants under the specific treatment tested, no other secondary metabolites are being produced that can compensate for the loss of melanin.

### Cell wall regulation

*E. dermatitidis* hosts a variety of chitin syntheses which are responsible for cell wall strengthening, cell wall remodelling, and protection from host immune systems (Abramczyk et al. 2009). All three conditional albinos had upregulation of multiple chitin synthase genes on YPG. A study conducted on *C. neoformans* demonstrated that mutant *chs3*Δ and *csr2*Δ strains both produced melanin but lost their ability to retain it (Banks et al. 2005). In *E. dermatitidis* the deletion mutant *wdchs1*Δ exhibited hyperpigmentation whereas the mutation *wdchs2*Δ did not have much impact on pigmentation (Zheng et al. 2006). In our study we saw upregulation of *CHS2* in the wildtype and all three conditional albinos but no upregulation of *CHS1* in either the wildtype or mutant strains. The study conducted by Zheng et al. (2006) also observed that addition of 1M sorbitol and temperature impacted the pigmentation in the *wdchs1*Δ and *wdchs2*Δ mutants but since our study was conducted at 28°C, the impact of these class 1 and class II synthase genes might not be that strong. In *E. dermatitidis* a class V chitin synthase (WdChs5p) is essential for sustained cell growth during infection and disruptions can lead to impaired melanin externalization affecting cell wall integrity (Liu et al. 2004). The upregulation of class V chitinase in the wildtype and the conditional albinos indicates the carbon source is most likely affecting cell wall organization in a manner that also impacts melanization.

### Metal acquisition

In fungi, iron plays an essential role in various processes and also acts as a cofactor for many different enzymes (Do et al. 2020). In pathogenic fungi, iron is a key nutrient for survival in the host environment where iron availability is limited. To overcome this, many pathogenic fungi have developed methods such as reductive iron transport and use of siderophore and heme uptake systems (Do et al. 2020; Voß et al. 2020). Siderophores are secondary metabolites that have a high affinity for ferric iron making them crucial for iron acquisition (Choi et al. 2024). In a study conducted by Chen et al. (2014), it was found that *E. dermatitidis* has multiple iron transport pathways and genes necessary to acquire iron from hosts during infection. This might also enable them to uptake iron from various niches hence providing an advantage against competitive species. In *E. dermatitidis* multiple enzymes responsible for the synthesis for siderophores were found upregulated during skin infection providing evidence of their importance during infection (Poynter et al. 2016; 2018). In our study genes responsible for iron acquisition were found to be significantly downregulated in the wildtype and all our albino strains.

Besides iron, copper also plays an important role in the production of melanin by participating in both the DHN and L-DOPA pathways (Walton et al. 2005; Saitoh et al. 2010; Eisenman and Casadevall 2015). We did not see any significant upregulation or downregulation of the copper transporter or copper chaperone genes indicating that the nutrient composition of the two treatments did not have a major influence on copper homeostasis.

### Carbon regulation

In fungi, D-galactose is catabolized via two different pathways: the Leloir pathway and oxido-reductive catabolic pathway (Fekete et al. 2004). Older studies looking at effects of galactose on the growth of fungi observed that galactose has an inhibitory effect on fungal growth (Horr 1936; Edgecombe 1938). In *E. dermatitidis,* upregulation of genes involved in the gluconeogensis pathway and the glyoxylate cycle have been observed during skin infection (Poynter et al. 2016). This includes the Snf1 kinase which is activated under gluconeogenic conditions leading to the activation of gluconeogensis (Poynter et al. 2016). Since production of DHN-melanin is controlled by the PKS1 pathway in *E. dermatitidis* specific carbon sources might be better suited to produce the starting materials or the intermediates. Different carbon sources may cause shifts in various pathways that might be involved in the production of L-DOPA-melanin or pyomelanin or they might also be better suited to upregulate the expression of genes involved in those pathways.

## Conclusion

The main purpose of this study was to perform RNA-seq analysis of the three conditional albino mutants (*alb1*, *alb2* and *alb3*) that had the ability to recover melanin production on different carbon sources even with mutations in *PKS1*. RNA-seq analysis demonstrated that the number of DEGs was much higher in the conditional *alb* mutants compared to the two obligate *alb* mutants. ORA analysis showed GO terms associated with cell membrane and transport were shared between all conditional albino mutants which might play a role in the cell wall deposition of melanin. There was upregulation of genes involved in the 1,8-DHN melanin and pyomelanin and various chitin synthase in all three conditional *alb* mutants indicating that melanization and cell wall regulation in fungi tend to work together. Further experiments can utilize knockout strategies to delete certain genes in the DOPA and pyomelanin pathways to observe the effects on melanization in the conditional *alb* mutants. Another approach can be to utilize IR spectra studies to determine if the melanin produced by the conditional *alb* mutants is similar to the melanin produced by the wildtype.

## Data availability

All data necessary for confirming the conclusions presented in this manuscript are present within the document, figures, and tables. Strains are available upon request. All RNA raw sequence reads (RNA-seq analysis) have been deposited under the BioProject accession: PRJNA1196592

## Supporting information

Supplemental File with GO term genes

## Supplemental Figures

**Figure S1:**
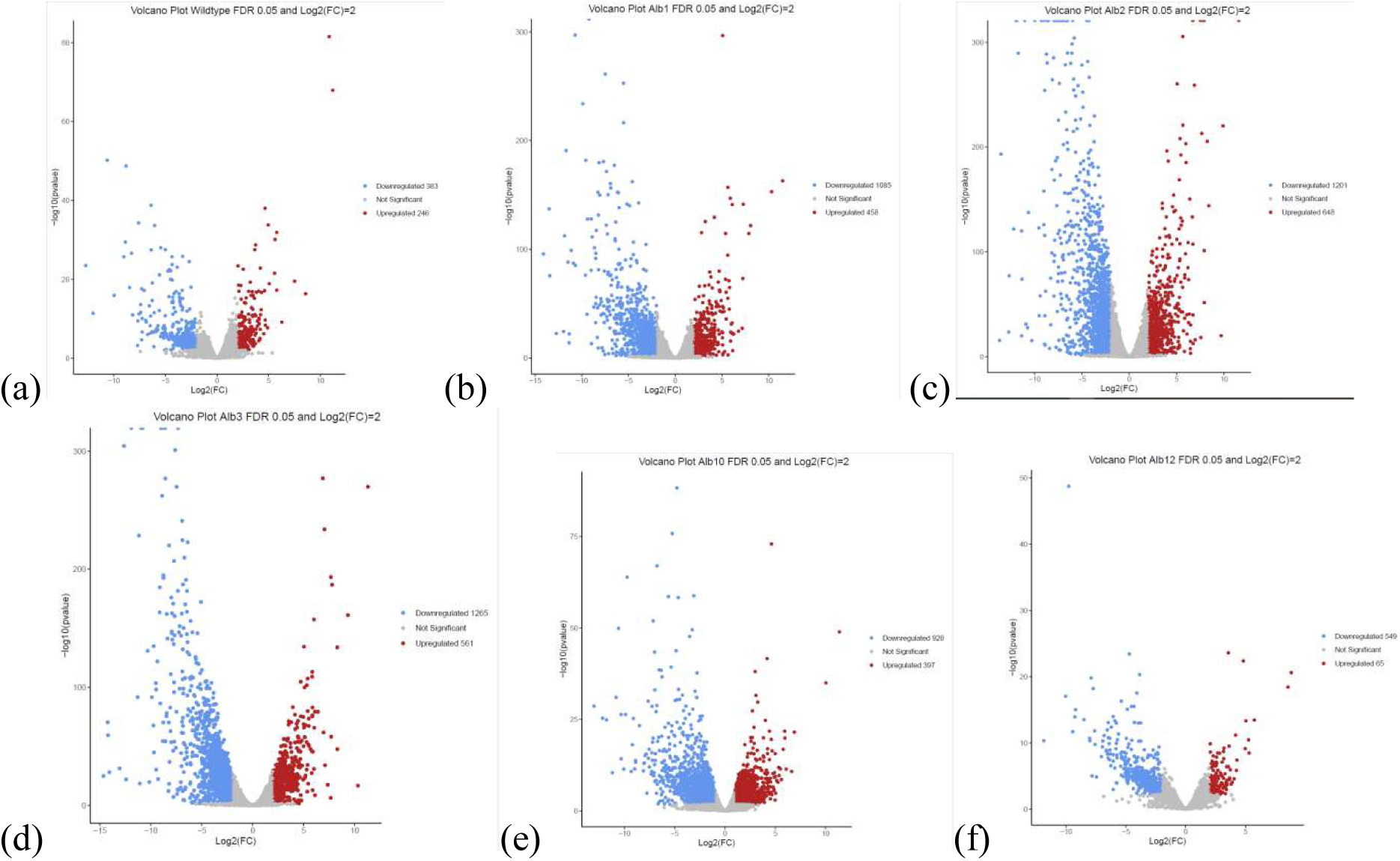
Volcano plots representing the number of differentially expressed upregulated (red dots) and downregulated (blue dots) genes observed on YPD vs YPG based on FDR <0.05 and Log2(FC) >2 or <-2 for (a) *E. dermatitidis* wildtype, (b) *alb1*, (c) *alb2*, (d) *alb3*, (e) *alb10*, and

**Figure S2:**
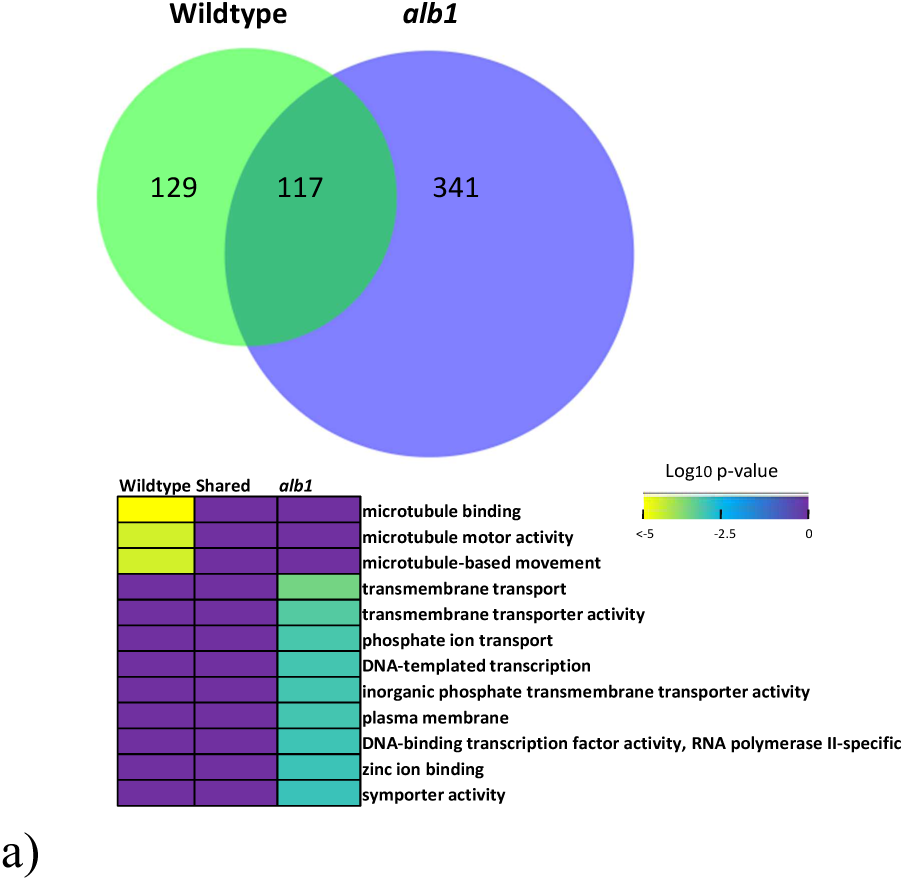
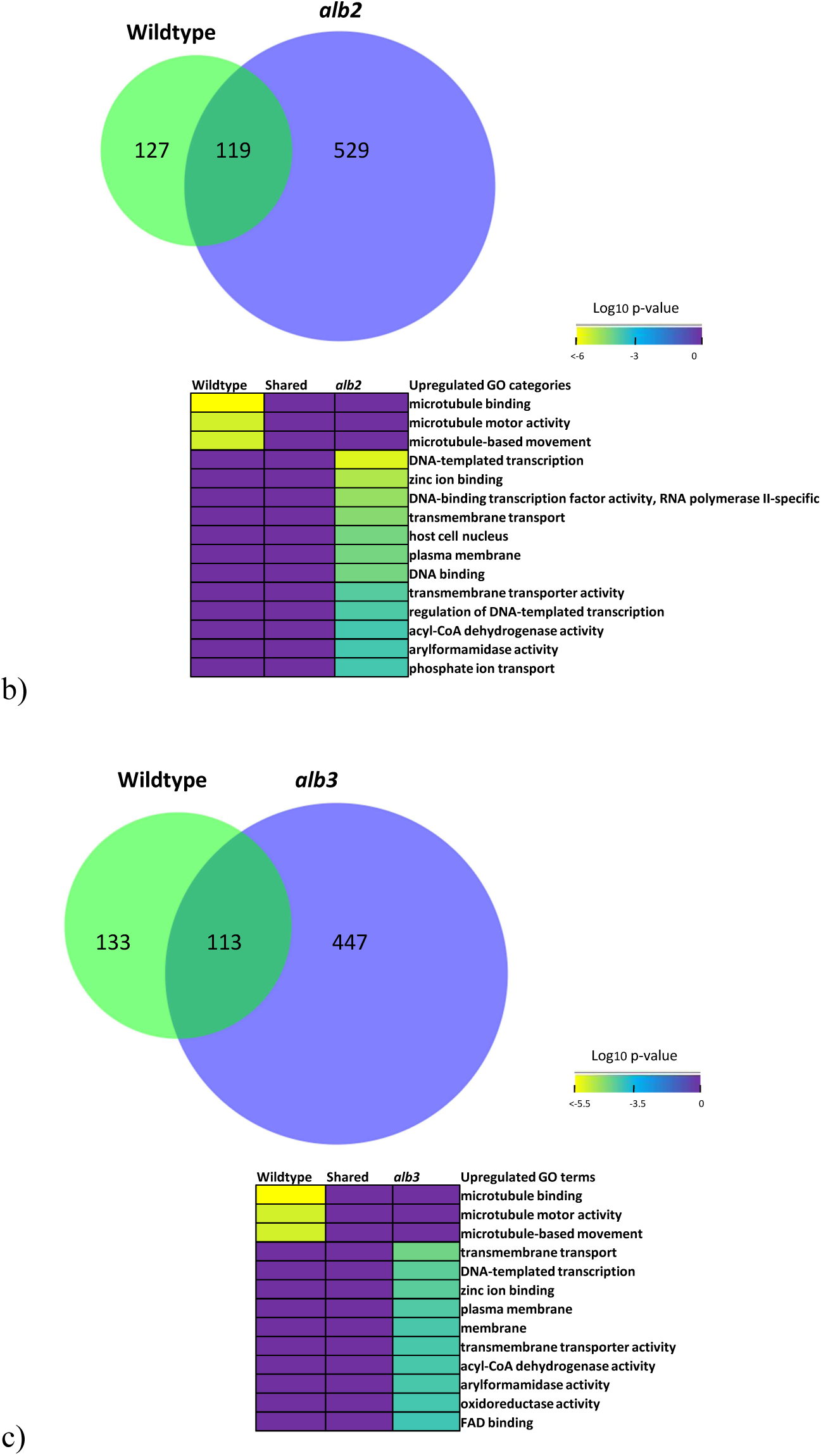
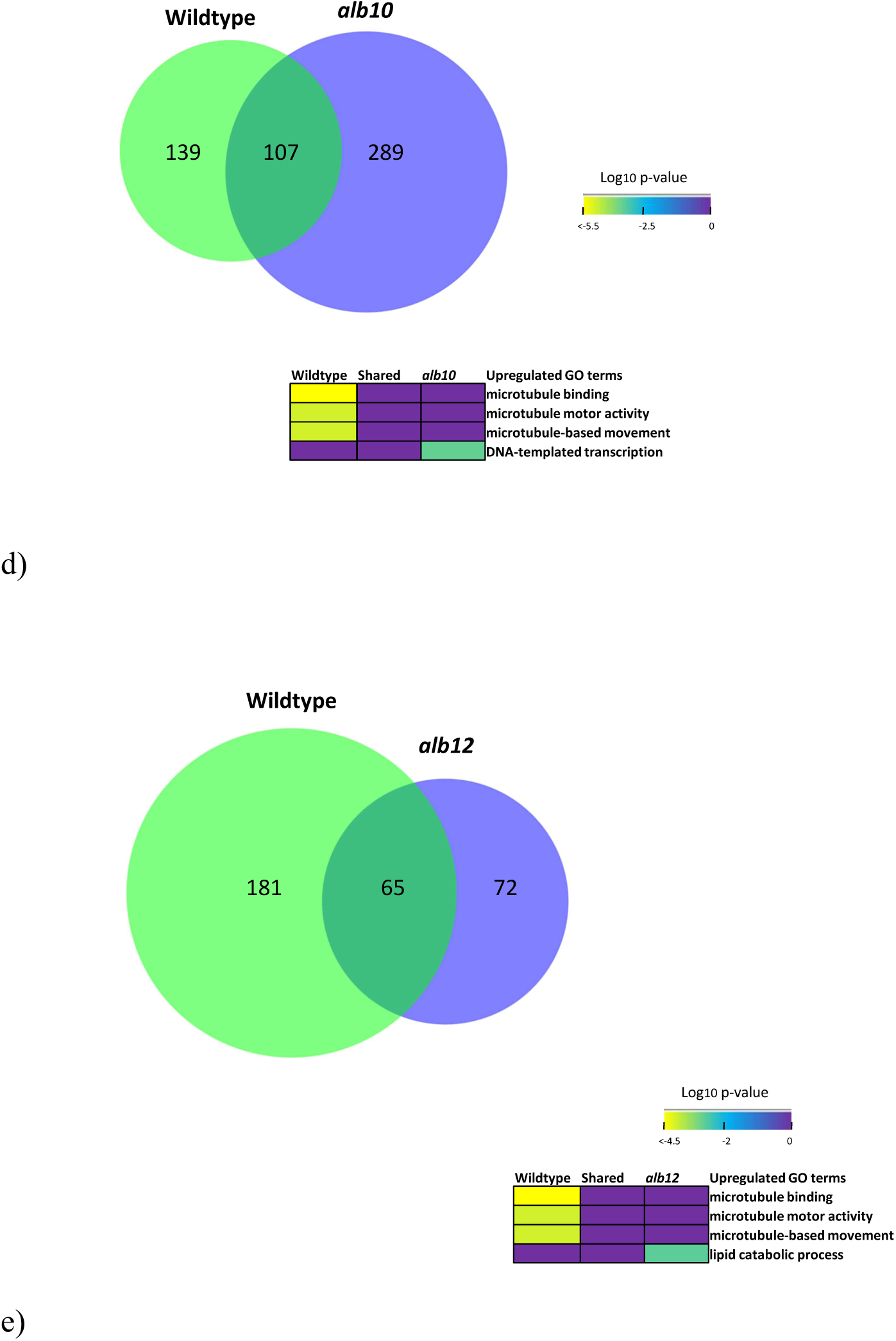
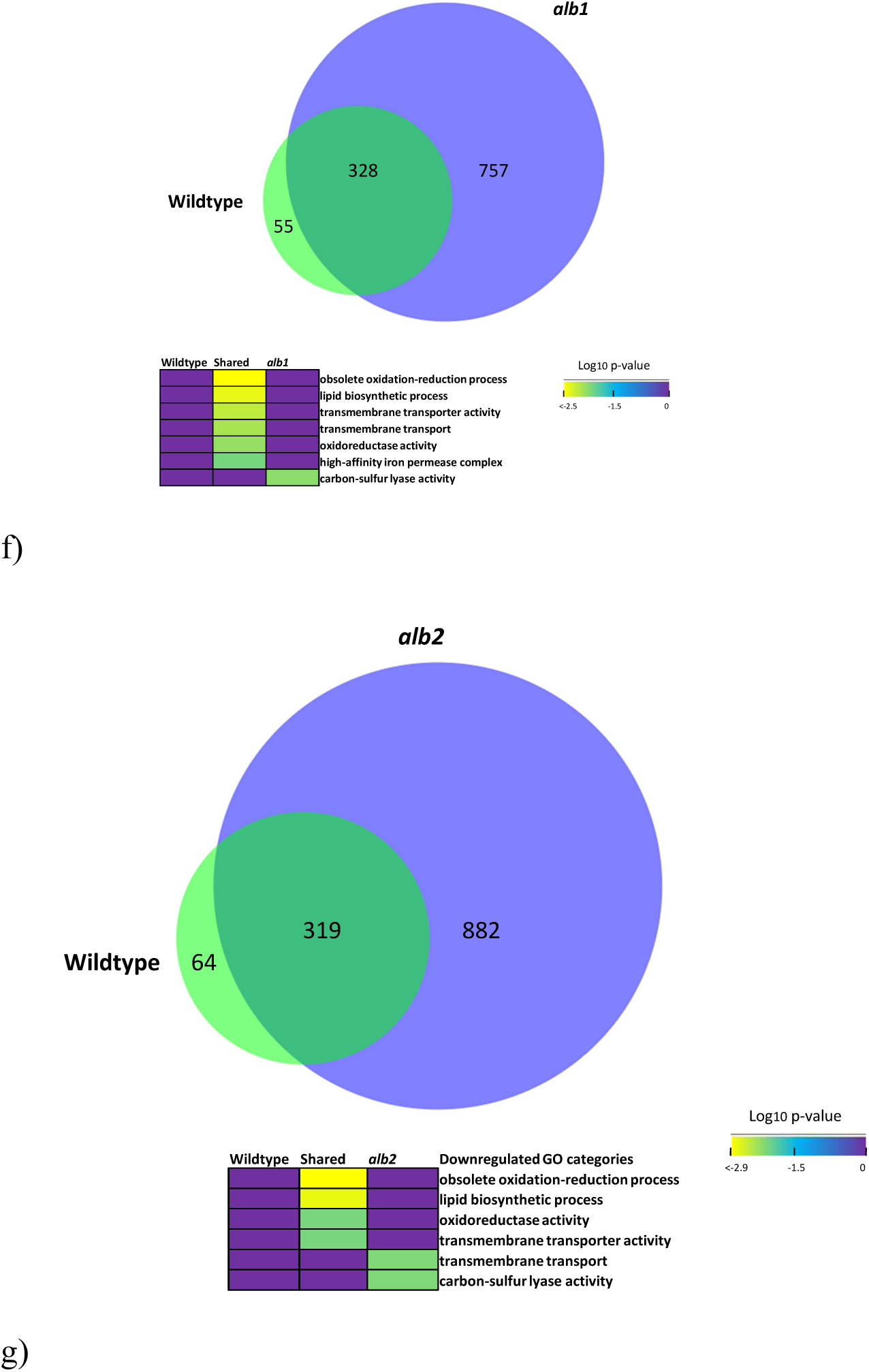
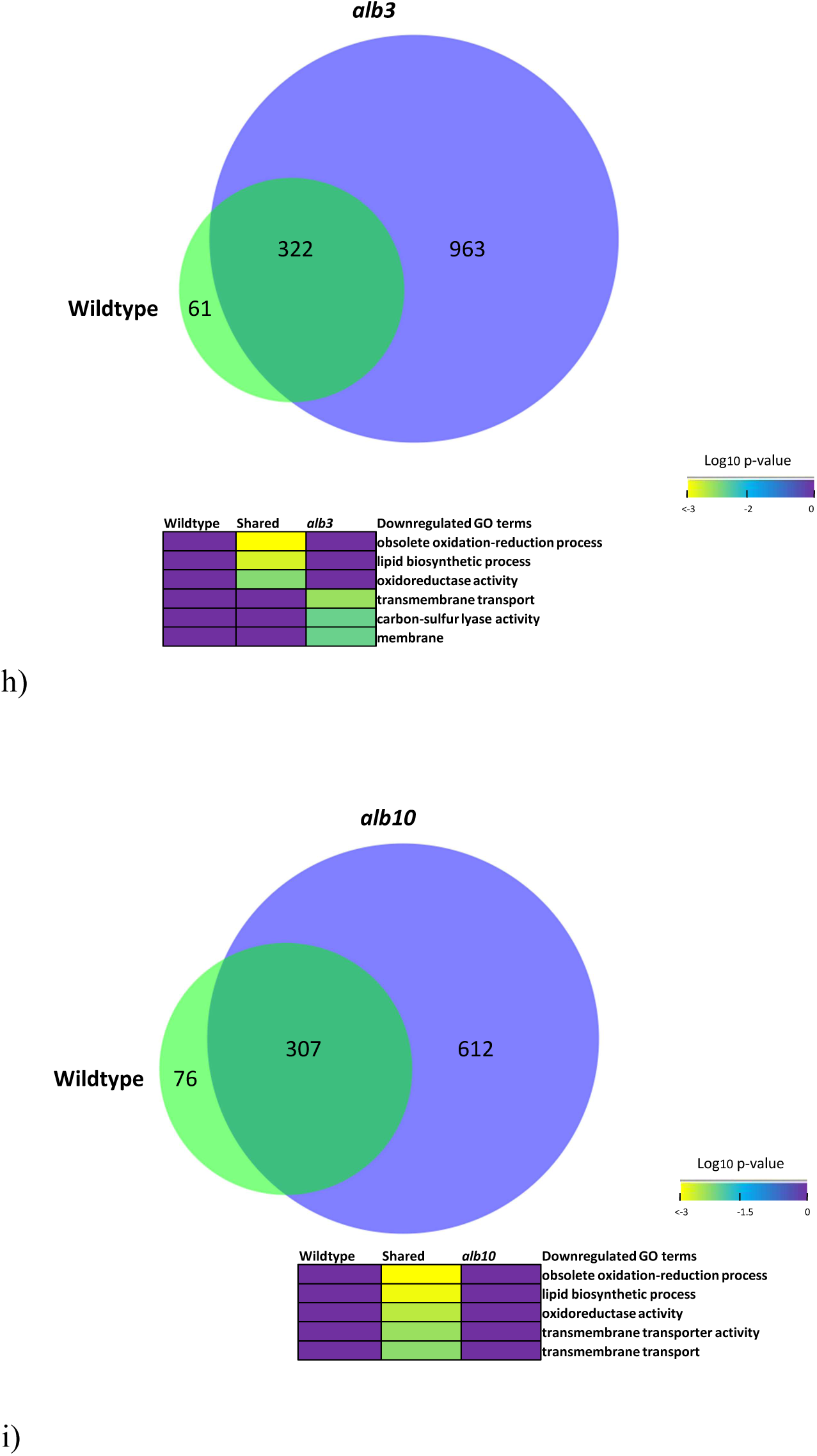
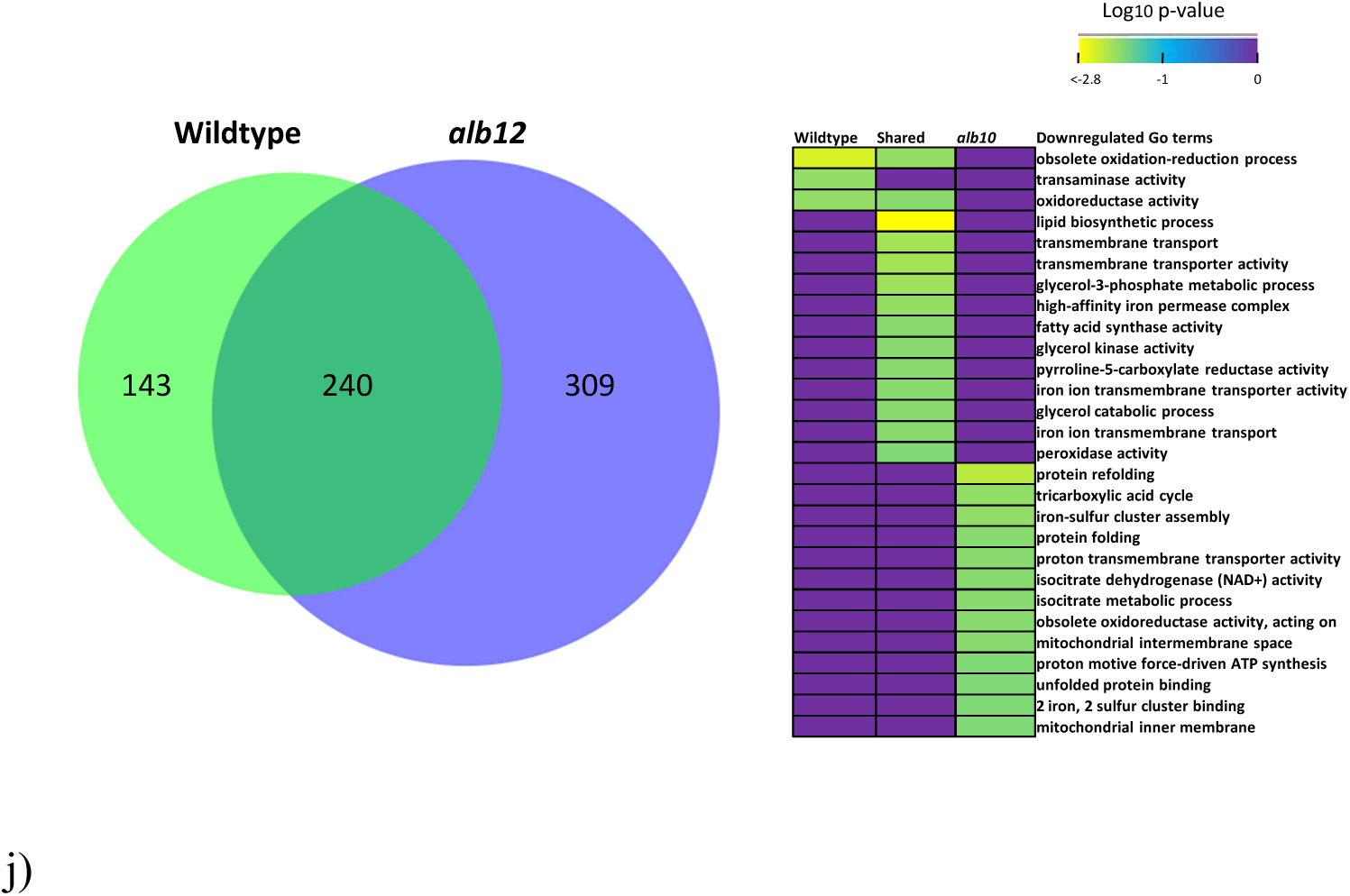
Venn diagram demonstrating the overlap in differentially expressed upregulated or downregulated genes in *E. dermatitidis* wildtype (WT) and albino mutant datasets. DEGs identified using DESeq2 with a significance cutoff of *p* < 0.05 and log_2_ FC > 2 or < −2 (a) WT and *alb1* upregulated, (b) WT and *alb2* upregulated, (c) WT and *alb3* upregulated, (d) WT and *alb10* upregulated, (e) WT and *alb12* upregulated, (f) WT and *alb1* downregulated, (g) WT and *alb2* downregulated, (h) WT and *alb3* downregulated, (i) WT and *alb10* downregulated, (j) WT and *alb12* downregulated. Terms presented for each section include Gene Ontology-terms that are considered significantly enriched at FDR < 0.05, from FungiFun 3.0. Yellow colour = more statistically enriched, purple colour= less statistically enriched.

